# Intrinsic timing, not temporal prediction, underlies ramping dynamics in visual and parietal cortex, during passive behavior

**DOI:** 10.1101/2025.09.11.673960

**Authors:** Yicong Huang, Ali Shamsnia, Mengze Chen, Shuang Wu, Timothy Stamm, Sophie Medico, Farzaneh Najafi

## Abstract

Ramping neural activity is widely interpreted as a signature of predictive processing, but whether such signals truly reflect predictions or instead emerge from sensory mechanisms remains unclear. To address this question, we used two-photon calcium imaging across multiple cell types in visual and parietal cortex while awake mice passively received repeated audiovisual stimuli presented under distinct temporal structures. Neurons segregated into two broad response classes: stimulus-activated (ramp-down) and stimulus-inhibited (ramp-up) populations with diverse temporal kinetics. Multiple findings argued against a predictive interpretation: ramping activity was present in naïve animals, neural responses changed immediately after short/long interval transitions, and unexpected stimulus timings elicited nearly identical responses in predictable and irregular contexts. Population analyses further showed that ramping reflected relaxation from stimulus-evoked activity rather than anticipatory buildup. Heterogeneous kinetics generated a robust population code for elapsed time. Together, these findings show that neural ramps during passive stimulation arise from stimulus-evoked dynamics that intrinsically generate temporal signals, rather than from temporal predictive processing.

**Teaser:** Ramping activity in visual-parietal cortex during passive stimulation arises from sensory responses, not predictive mechanisms.

## INTRODUCTION

The brain continuously encodes the passage of time and generates expectations about when future events will occur. These temporal computations are critical for perception, motor control, and learning, yet their neural underpinnings remain debated (*1*).

Neural activity between sensory events may reflect two distinct processes: elapsed-time signaling, which encodes the duration since the previous event, and temporal predictive processing, which represents the expected timing of the next event. These processes unfold concurrently; hence, it is often difficult to determine whether neural responses in a given circuit reflect temporal tracking or predictive processing mechanisms.

### Temporal signaling

the encoding of elapsed time, is achieved through multiple mechanisms (*2, 3*), such as ramping (gradual changes in neural activity) (*4-6*), sequential activation of neurons (*7*), and population trajectories, where activity patterns of recurrently-connected neurons evolve through a high-dimensional state space, with each state carrying information about elapsed time (*8-12*).

Synaptic mechanisms, such as short-term plasticity that modulate how neurons respond to inputs over time (*13, 14*), together with variability in intrinsic time constants, generate heterogeneous dynamics across the population. This diversity acts as a temporal basis set: overlapping dynamics tile different segments of an interval, allowing downstream circuits to reconstruct elapsed time from distributed activity. Such population-level coding has been observed across cortical, striatal, hippocampal, and cerebellar circuits (*15-24*).

In the cortex, both sensory and associative areas demonstrate temporal processing. For example, neurons in the posterior parietal cortex (PPC) contribute to interval production and categorization (*25-29*), and neurons in frontal/prefrontal cortices demonstrate population trajectories that flexibly “scale” with the demanded interval length (*9, 30, 31*). Primary sensory cortices also show timing signatures when stimuli are predictable: neurons in primary visual and auditory cortex “remember” recent intervals, through stimulus-evoked state shifts that depend on short-term depression/facilitation (*32-34*).

### Predictive processing

on the other hand, suggests that cortical areas contain internal models of the world, which generate predictions that are compared with sensory inputs. When a mismatch occurs, prediction error (PE) signals are generated to update the internal model (*35-38*). This canonical computation has been identified across many brain regions and species, and applies to both the identity and the timing of sensory inputs— that is, *what* and *when* a stimulus will occur (*38*). Multiple cortical areas—including visual and parietal cortex—have been implicated in predictive processing (*38-46*).

### Cell-type-specific

studies demonstrate a key role for inhibitory neurons— particularly VIP and SST interneurons—in predictive processing (VIP: vasoactive intestinal peptide; SST: somatostatin). VIP neurons in primary and higher visual areas respond to the change (*44, 47*) or omission (*41, 43, 44, 48-50*) of expected visual stimuli.

SST interneurons, in the PPC, include a molecularly defined subset that carries an error-correction signal during navigation tasks (*51*). Computational modelling and circuit dissection suggest that VIP–SST–Excitatory disinhibitory circuit can give rise to prediction-error neurons (*52, 53*).

In temporal processing, too, inhibitory neurons play a role: SST interneurons are required for sequential population activity in both motor (*54*) and prefrontal cortex (*55*). Particularly relevant to the present study is the ramping activity—observed in most VIP and some excitatory neurons—prior to an *expected* stimulus. Importantly, this ramp continues even further, when that stimulus is omitted (*41-43*).

### Disambiguating temporal signaling from predictive processing

Temporal processing and temporal predictive processing can co-occur; however, their computational requirements differ. Predictive processing relies on learned internal models and therefore suggests that prediction signals should strengthen with experience. Temporal signaling, in contrast, can arise intrinsically from recurrent or synaptic dynamics and should be present even in naïve animals exposed to irregular stimulus statistics.

Many experimental paradigms entangle these processes. Regularly repeated stimuli simultaneously provide temporal signals and high predictability, making it difficult to attribute neural responses—such as ramping before a predictable event or activity after an unexpected sensory omission—to one mechanism or the other.

Disambiguating these mechanisms is crucial for identifying the true computational role of neural circuits and for distinguishing intrinsic dynamics from learned predictive processing.

### Present Study

We devised a three-stage paradigm in awake, head-fixed mice to tease apart temporal predictive processing and temporal signaling while performing two-photon calcium imaging of layer 2/3 excitatory, VIP, and SST neurons in medial visual cortex (VIS) and PPC. Across all experiments, mice passively viewed repeating audio-visual stimuli (0.2 s), separated by inter-stimulus intervals (ISIs)—neutral gray screen and no sound—whose durations varied according to the paradigm design.

1. <u>Random ISI sessions:</u> naïve mice received stimuli at uniformly distributed ISIs (0.5– 2.5 s). High temporal variability minimizes timing predictions. Interval-dependent activity is therefore primarily attributed to temporal processing.
2. <u>Fixed/Jittered ISI sessions:</u> blocks of trials with fixed ISI (1.5 s)—hence fully predictable—alternated with blocks of jittered ISIs (0.5-2.5 s). Both blocks included occasional deviant ISIs (0.5 or 2.5 s: shorter or longer than expected), representing temporal prediction errors. Predictive processing predicts weaker anticipatory and prediction-error signals under jitter, whereas temporal processing should encode interval length regardless of variability.
3. <u>Short/Long ISI blocks:</u> mice experienced trial blocks alternating between 1-s or 2-s fixed ISIs, allowing us to test whether block transitions elicited prediction-error responses, and whether new temporal predictions emerged within each block.

### Results summary

Neurons clustered into two major categories: stimulus-activated (ramp-down) and stimulus-inhibited (ramp-up). Stimulus-inhibited neurons exhibited gradual pre-stimulus increases in activity and are therefore referred to as ramp-up clusters. Conversely, stimulus-activated neurons showed decreasing activity leading up to stimulus onset and are referred to as ramp-down clusters. Each cluster contained functional subtypes defined by distinct rise and decay kinetics.

**(1)** Across all clusters, neural activity shifted immediately at transitions between short and long-ISI blocks, inconsistent with gradual prediction updating. Deviant intervals in fixed versus jittered-ISI blocks elicited no differential responses, and the same functional clusters were present even in naïve animals exposed to random intervals. **(2)** Population trajectories showed that ramping during stimulus intervals simply reflects the relaxation (recovery) of activity from the stimulus evoked state. **(3)** Decoding analysis revealed that heterogeneous neuronal kinetics give rise to a distributed population code for elapsed time.

Together, across multiple complementary experimental paradigms, our results argue that ramping activity in VIS/PPC during passive stimulation does not reflect temporal predictive processing, but instead arises from intrinsic stimulus-evoked dynamics that nonetheless support accurate decoding of elapsed time.

### Significance

These results resolve longstanding ambiguities by showing that ramping dynamics—previously interpreted as predictive signals in visual cortex—instead reflect stimulus-evoked activity whose intrinsic evolution encodes elapsed time (*56*). More broadly, they underscore the importance of incorporating carefully designed jittered paradigms before attributing ramping or omission-related activity to prediction or prediction-error signals. Key next steps are to: **(1)** identify the circuits that implement temporal predictive processing during passive perception; **(2)** determine whether, in perceptual decision tasks, VIS/PPC dynamics remain signatures of elapsed time or shift toward predictive coding; and **(3)** dissect the cellular and synaptic mechanisms underlying intrinsic temporal codes in cortex.

## RESULTS

### Paradigms dissociating temporal predictive processing from stimulus-evoked temporal dynamics

Neural signals during the interval between two events could reflect either of two processes (**Fig. 1A**): **(1)** the elapsed time since the previous event (temporal signaling), or **(2)** the expected timing of the next event (temporal predictive processing). These processes unfold concurrently, making it difficult to determine which function a neural circuit represents. Disentangling them is essential, because they rely on distinct underlying computations, and failing to do so risks misinterpreting circuit function.

**Figure 1.**
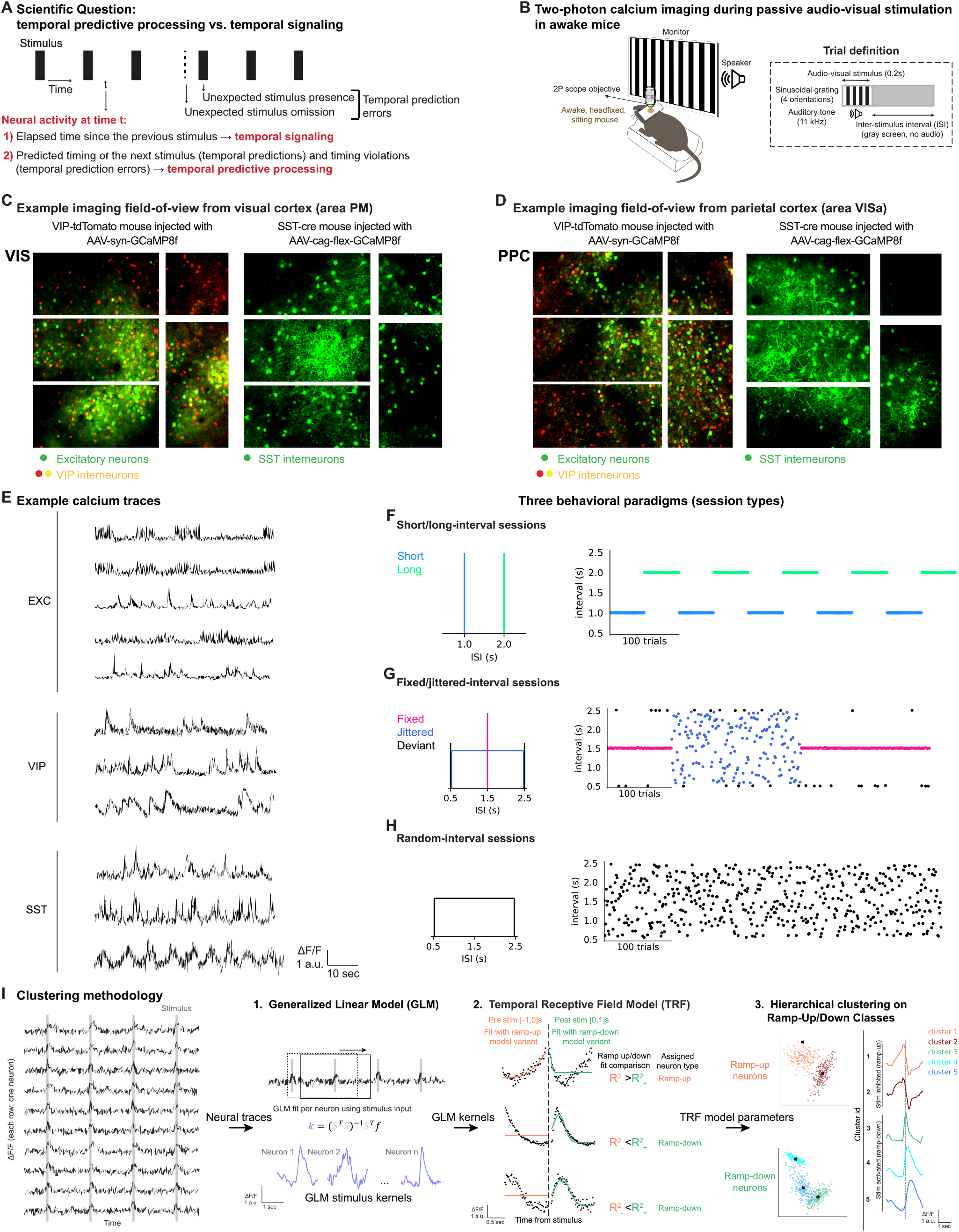
Experimental paradigms dissociating temporal predictive processing from stimulus-evoked temporal dynamics. **(A)** Neural activity at each timepoint could reflect one of the 2 processes: **(1)** temporal signaling, i.e., encoding elapsed time since the previous stimulus; or **(2)** Temporal predictive processing, i.e., generating predictions and prediction errors about the timing of the upcoming stimulus. Temporal prediction errors occur when a stimulus is presented earlier or later (‘omission’) than expected. **(B) Left:** Awake, head-fixed mice received repeating audio-visual stimuli (0.2s), separated by an inter-stimulus interval (ISI; gray screen), while two-photon calcium imaging was performed from the medial visual cortex (area PM) and posterior parietal cortex (PPC, area VISa). **Right:** A trial is defined as a 200ms audio-visual stimulus followed by an ISI with a gray screen and no auditory tone. ISI duration varied across paradigms (see F-H). **(C)** Visual cortex imaging field-of-view. Two example sessions. **Left:** VIP-tdTomato mouse injected with GCaMP8f. All neurons express GCaMP8f; a subset is in red, labeling VIP interneurons. (Red and yellow: VIP interneurons. Green: non-VIP neurons, i.e., presumptive excitatory neurons). **Right:** SST-cre mouse injected with cre-dependent GCaMP8f. (Green: SST interneurons). **(D)** Same as C, but for PPC. **Fig. S1** shows histological analysis of the imaged sites. **(E)** Example fluorescence traces for excitatory, VIP, and SST neurons. **(F–H)** Mice received three paradigms (session types) distinguished by their ISIs. **Left:** distribution of ISIs in a session. **Right:** ISIs for example consecutive trials in a session. **(F)** Short/long sessions had alternating blocks of trials with short (1s; blue) and long (2s; green) ISIs. (**G)** Fixed/jittered sessions had alternating blocks of trials with fixed (1.5s; red) and jittered (0.5–2.5s; blue) ISIs. Each block also included 4% deviant ISIs (0.5s and 2.5s, black). (**H)** In random sessions all trial ISIs were randomly chosen from a uniform distribution (0.5–2.5s). **(I)** Clustering methodology. (**1)** <u>GLM model:</u>neural responses across the full session were modeled with a GLM using stimuli as input. (**2)**<u>TRF model:</u> two temporal-receptive-field model variants, with opposite input polarity, were applied to the pre- and post-stimulus components of each kernel to get the ramp-up and ramp-down model responses, respectively. Goodness-of-fit comparisons determined whether each neuron was classified as ramp-up or ramp-down. (**3)** <u>Hierarchical clustering:</u> finally, neurons within each class were clustered using K-means on the TRF model parameters.

To address this, we designed three paradigms to separate temporal signaling from predictive processing. Awake, head-fixed mice passively received a repeating audio-visual stimulus (0.2 s), separated by an inter-stimulus interval (ISI; 0.5–2.5 s; **Fig. 1B**; ISI includes gray screen and no auditory stimulus). Each stimulus consisted of a simultaneous auditory tone (11 kHz) and visual grating (whole-field static black/white stripes). Two-photon calcium imaging, via GCaMP8f (*57*), was performed from the medial visual cortex (VIS; area PM, also termed V2M) and the posterior parietal cortex (PPC; area VISa; **Fig. S1**). Layer 2/3 excitatory neurons and VIP interneurons were simultaneously imaged in one cohort, and SST interneurons were imaged in a separate cohort of mice (VIP-Cre;Ai14 mice and SST-Cre mice injected with GCaMP8f; **Fig. 1C-E**).

The three paradigms were short/long, fixed/jittered, and random sessions (**Fig. 1F-H**).

- In short/long sessions, two block types alternated (50 trials each): ISI was fixed at 1 s in short blocks and 2 s in long blocks (**Fig. 1F**).
- In fixed/jittered sessions, two block types alternated (200 trials each): ISI was fixed at 1.5 s in fixed blocks, and randomly drawn from a uniform 0.5–2.5 s distribution in jittered blocks (**Fig. 1G**). Each block also contained 4% deviant ISIs (short deviant: 0.5 s; long deviant: 2.5 s). Deviants carried high surprisal in fixed blocks, while in jittered blocks they lay within the expected ISI range.
- In random sessions, all ISIs were independently drawn from a uniform distribution (0.5–2.5 s; **Fig. 1H**).

Importantly, naïve mice were first exposed to the random sessions, before experiencing the short/long and fixed/jittered paradigms. This design allowed us to examine neural responses to random intervals in animals without prior exposure to our stimuli.

### VIS and PPC neurons exhibit stimulus-activated and pre-stimulus ramping responses

We measured the response of excitatory, VIP, and SST neurons in VIS and PPC during passive audio-visual stimulation, using two-photon calcium imaging (**Table 1**). The same field-of-view was imaged across days to enable direct comparison of neural responses across the three paradigms. Neurons in both areas showed diverse response profiles (**Fig. 2A,G**), hence we clustered them into functional subtypes (**Fig. 1I**).

**Table 1.** Number of total imaged neurons per brain area, cell type, and paradigm.

| Region | Cell type | Random | Fixed/jittered | Short/long | Total |
| --- | --- | --- | --- | --- | --- |
| VIS | Excitatory | 2,226 | 5,634 | 7,101 | 14,961 |
|  | VIP | 597 | 692 | 1,233 | 2522 |
|  | SST | 181 | 600 | 640 | 1421 |
|  | Total | 3,004 | 6,926 | 8,974 | 18,904 |
| PPC | Excitatory | 2,437 | 5,327 | 3,913 | 11,677 |
|  | VIP | 304 | 678 | 401 | 1,383 |
|  | SST | 319 | 839 | 617 | 1,775 |
|  | Total | 3,060 | 6,844 | 4,931 | 14,835 |

**Figure 2.**
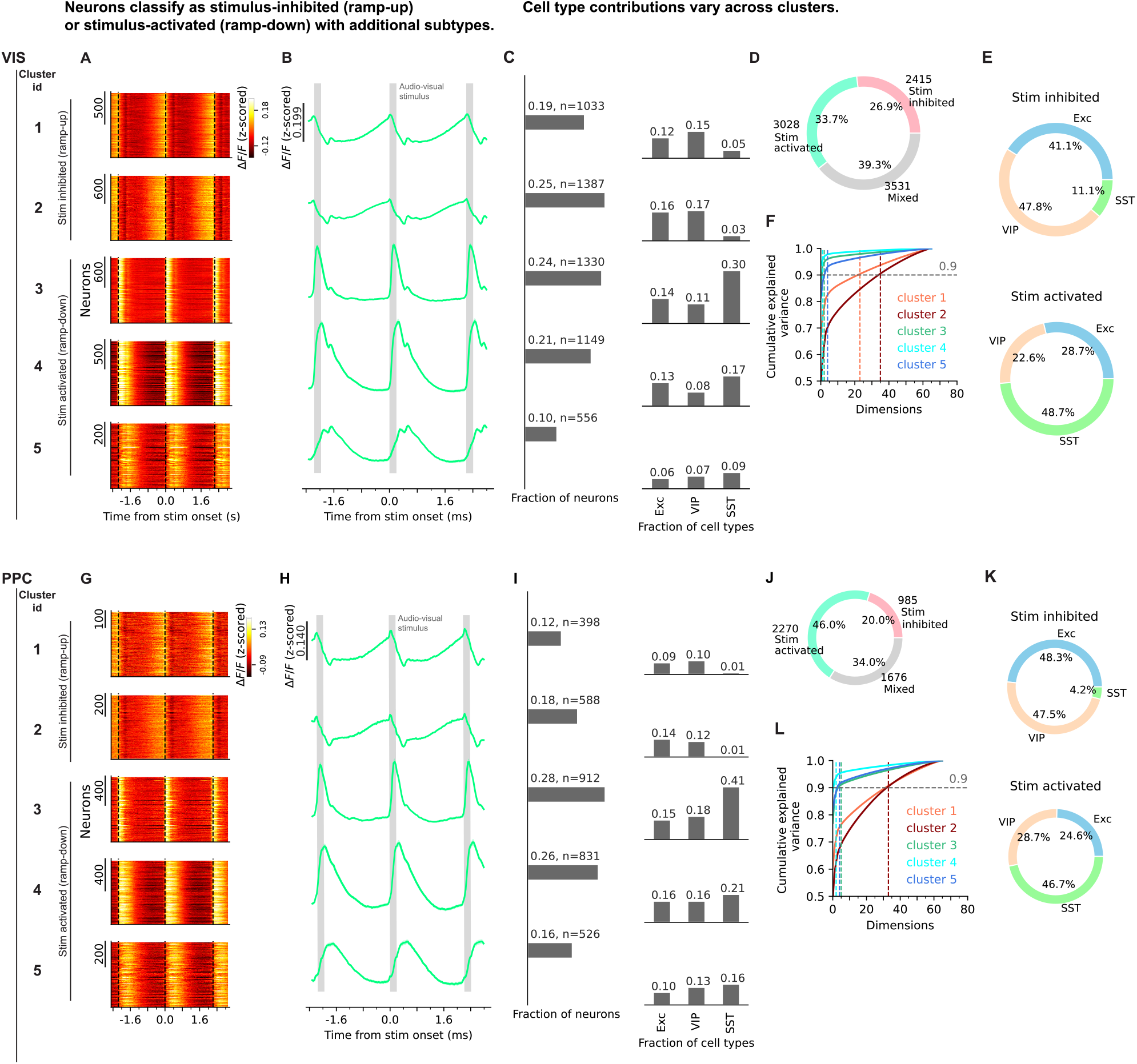
VIS and PPC neurons demonstrate stimulus-evoked responses, and pre-stimulus ramping activity to repeating audio-visual stimuli. Data from the long block of short/long sessions. Clusters were identified on pooled neurons across all short/long sessions. **VIS:** top panels; **PPC:** bottom panels. **(A)** Heatmaps show trial-averaged, single-neuron activity (ΔF/F) for each cluster (top to bottom). The stimulus-inhibited (ramp-up) and stimulus-activated (ramp-down) labels on the left are based on the temporal-receptive-field (TRF) model outcome; the subtypes within each group (heatmap) are identified by k-means clustering applied on TRF model parameters (**Fig. 1I**). A 3^rd^ class of neurons with mixed ramp-up and ramp-down dynamics (‘mixed’ cluster) is excluded from the main Figures, but illustrated in detail in **Fig. S3, Table 2,3**, and (**D)** here. **(B)** ΔF/F across neurons and trials, for each cluster (top to bottom). Mean ± 95% confidence interval, across pooled neurons and trials from all sessions of all mice. Gray boxes: stimuli. **(C) Left:** fraction of all neurons assigned to each cluster. **Right:** fraction of neurons of each cell type assigned to each cluster. **(D)** Fraction of neurons belonging to the two broad response classes, along with the “mixed” cluster, which did not meet the TRF-model criteria for ramp-up or ramp-down classification. Fractions shown in panels **(C)** and **(D)** exclude the mixed cluster. **(E)** Fraction of genetically-defined cell types within each broad class of stimulus-inhibited (top), and stimulus-activated (bottom) neurons. To account for differences in the number of neurons across genetic cell types (excitatory, VIP, SST), each functional subtype was first normalized to the total number of neurons within its corresponding genetic class. **(F)** PCA performed on trial-averaged population activity traces (one stimulus and one ISI) for each functional cluster. Cumulative explained variance as a function of population dimensionality. Colored lines indicate the number of dimensions required to explain 90% of the variance for each cluster. **(G-L)** Same as **(A-F)**, but for PPC.

To identify response classes, we first fit single-neuron responses in each session using a generalized linear model (GLM) with a single stimulus kernel (**Fig. 1I,1**). We then fit each neuron’s GLM kernel using an exponentially modified Gaussian temporal receptive field (TRF) model (*58, 59*). The TRF model classified neurons into two broad response classes: ramp-up and ramp-down (**Fig. 1I,2**), and yielded parameters describing response dynamics, including ramp speed and temporal width (**Fig. 7E,K**; **Fig. S2,10**). Hierarchical clustering of these TRF-derived parameters then identified functional subtypes within each broad class (**Fig. 1I,3**).

**Table 2.** Number (fraction) of imaged neurons in the visual cortex (VIS) classified into functional clusters, across cell types and paradigms. Fractions represent the proportion of neurons within a given category relative to the total number of neurons— across stimulus-inhibited, stimulus-activated, and mixed clusters—within the same brain area, cell type, and paradigm. For example, the top-left cell indicates that 17% of excitatory neurons recorded in VIS during random sessions belonged to the stimulus-inhibited cluster.

|  |  | **Random** | **Fixed/jittered** | **Short/long** | **Extended-ISI**<br>**random** |
| --- | --- | --- | --- | --- | --- |
| **Excitatory** | **Stim-inhibited** | 387 (0.17) | 1358 (0.24) | 884 (0.23) | 800 (0.43) |
|  | **Stim-activated** | 913 (0.41) | 2113 (0.38) | 1600 (0.41) | 548 (0.23) |
|  | **Mixed** | 926 (0.42) | 2163 (0.38) | 1429 (0.37) | 992 (0.42) |
| **VIP** | **Stim-inhibited** | 136 (0.23) | 163 (0.24) | 89 (0.22) | 300 (0.42) |
|  | **Stim-activated** | 177 (0.30) | 212 (0.31) | 191 (0.48) | 81 (0.11) |
|  | **Mixed** | 284 (0.48) | 317 (0.46) | 121 (0.30) | 332 (0.47) |
| **SST** | **Stim-inhibited** | 8 (0.04) | 40 (0.07) | 12 (0.02) | 0 (0.00) |
|  | **Stim-activated** | 98 (0.54) | 383 (0.64) | 479 (0.78) | 0 (0.00) |
|  | **Mixed** | 75 (0.41) | 177 (0.29) | 126 (0.20) | 0 (0.00) |

**Table 3.** Number (fraction) of imaged neurons in the PPC classified into functional clusters, across cell types and paradigms. Same as Table 2, but for PPC.

|  |  | **Random** | **Fixed/jittered** | **Short/long** |
| --- | --- | --- | --- | --- |
| **Excitatory** | **Stim-inhibited** | 166 (0.07) | 829 (0.16) | 1969 (0.28) |
|  | **Stim-activated** | 1589 (0.65) | 2921 (0.55) | 2348 (0.33) |
|  | **Mixed** | 682 (0.28) | 1577 (0.30) | 2784 (0.39) |
| **VIP** | **Stim-inhibited** | 9 (0.03) | 96 (0.14) | 398 (0.32) |
|  | **Stim-activated** | 242 (0.80) | 393 (0.58) | 321 (0.26) |
|  | **Mixed** | 53 (0.17) | 189 (0.28) | 514 (0.42) |
| **SST** | **Stim-inhibited** | 6 (0.02) | 4 (0.00) | 48 (0.07) |
|  | **Stim-activated** | 253 (0.79) | 690 (0.82) | 359 (0.56) |
|  | **Mixed** | 60 (0.19) | 145 (0.17) | 233 (0.36) |

Using this approach, VIS and PPC neurons separated into two major response classes: stimulus-inhibited neurons with pre-stimulus ramp-up dynamics, and stimulus-activated neurons with ramp-down dynamics (**Fig. 2**; **Table 2,3**). Each class contained additional subtypes with distinct temporal kinetics. 37% of neurons—across both brain regions—were not strictly classified as ramp-up or ramp-down by the TRF model (“mixed” neurons; **Fig. 2D,J**; **Table 2,3**). These neurons led to the same conclusions as ramp-up/down populations (**Fig. S3**), and were excluded from the main figures for clarity (see Discussion).

Stimulus-inhibited responses evolved into a gradual ramp-up (**Fig. 2**; rows 1-2), whereas stimulus-activated responses evolved into a gradual ramp-down (**Fig. 2**; rows 3-5). Accordingly, we also refer to these clusters as ramp-up and ramp-down neurons, respectively. Both clusters contained subtypes with distinct response dynamics: stimulus-activated neurons ranged from fast/narrow to slow/broad responses; similarly, stimulus-inhibited neurons differed in ramp onset and speed (**Fig. 2**).

A secondary peak was visible in several population averages in both VIS and PPC (**Fig. 2B,H**). This peak occurred approximately 250 ms after stimulus offset. This timing is consistent with post-stimulus 2–10 Hz membrane-potential fluctuations reported in the visual cortex of awake stationary mice (*60*), and with state-dependent 3–5 Hz thalamocortical rhythms linked to movement and arousal offset (*61*).

Stimulus-inhibited ramp-up clusters were more frequent in VIS (44% VIS neurons vs. 30% PPC neurons; **Fig. 2C,I** left, rows 1-2). Stimulus-activated clusters with slow/broad peaks were more frequent in the PPC (31% VIS neurons vs. 42% PPC neurons; **Fig. 2C,I** left, rows 4-5). The prevalence of slow stimulus-driven clusters in PPC aligns with the role of higher-order associative areas in perception (*62*). The presence of ramp-up clusters in PPC confirms and extends previous reports of ramping neurons in V1 and higher visual area LM (*41, 43*).

Excitatory and inhibitory subtypes contributed to all clusters, but to varying degrees. SST neurons were most prevalent in stimulus-activated clusters (*42*) (**Fig. 2E,K** bottom). In contrast, a substantial fraction of VIP and excitatory neurons was stimulus-inhibited (*42*) (**Fig. 2E,K** top).

Population dimensionality differed systematically across clusters (**Fig. 2F,L**), with stimulus-activated neurons occupying substantially lower-dimensional subspaces than stimulus-inhibited (ramp-up) clusters (∼3 vs ∼50 dimensions for 90% explained variance). Overall, dimensionality was higher for clusters with slower and broader response profiles (compare **Fig. 2F,L** with **Fig. 7E,K**).

Population dimensionality in visual cortex is known to depend strongly on stimulus complexity and the animal’s behavioral state (*63, 64*). The lower dimensionality observed here is therefore consistent with our use of a restricted stimulus set (pure tones paired with four static gratings) and recordings in head-fixed mice on a stationary platform without overt locomotion. In contrast, studies using large sets of natural images in locomoting animals have reported substantially higher-dimensional population responses, reflecting more distributed and heterogeneous neural activity (*63*).

### Stimulus-activated and stimulus-inhibited responses are larger following longer intervals

To study how VIS/PPC responses encode distinct intervals, we compared neural activity between short- and long-ISI blocks in the short/long paradigm (**Fig. 1F**; short ISI: 1 s; long ISI: 2 s).

All clusters showed distinct activity patterns between the two blocks (**Fig. 3**; blue: short block; green: long block). Both response magnitude and ramp kinematics differed across short/long blocks (**Fig. 3B**). Stimulus-evoked response magnitudes —whether activated or inhibited— were smaller in the short block, potentially reflecting incomplete recovery from adaptation at shorter ISIs (*65, 66*).

**Figure 3.**
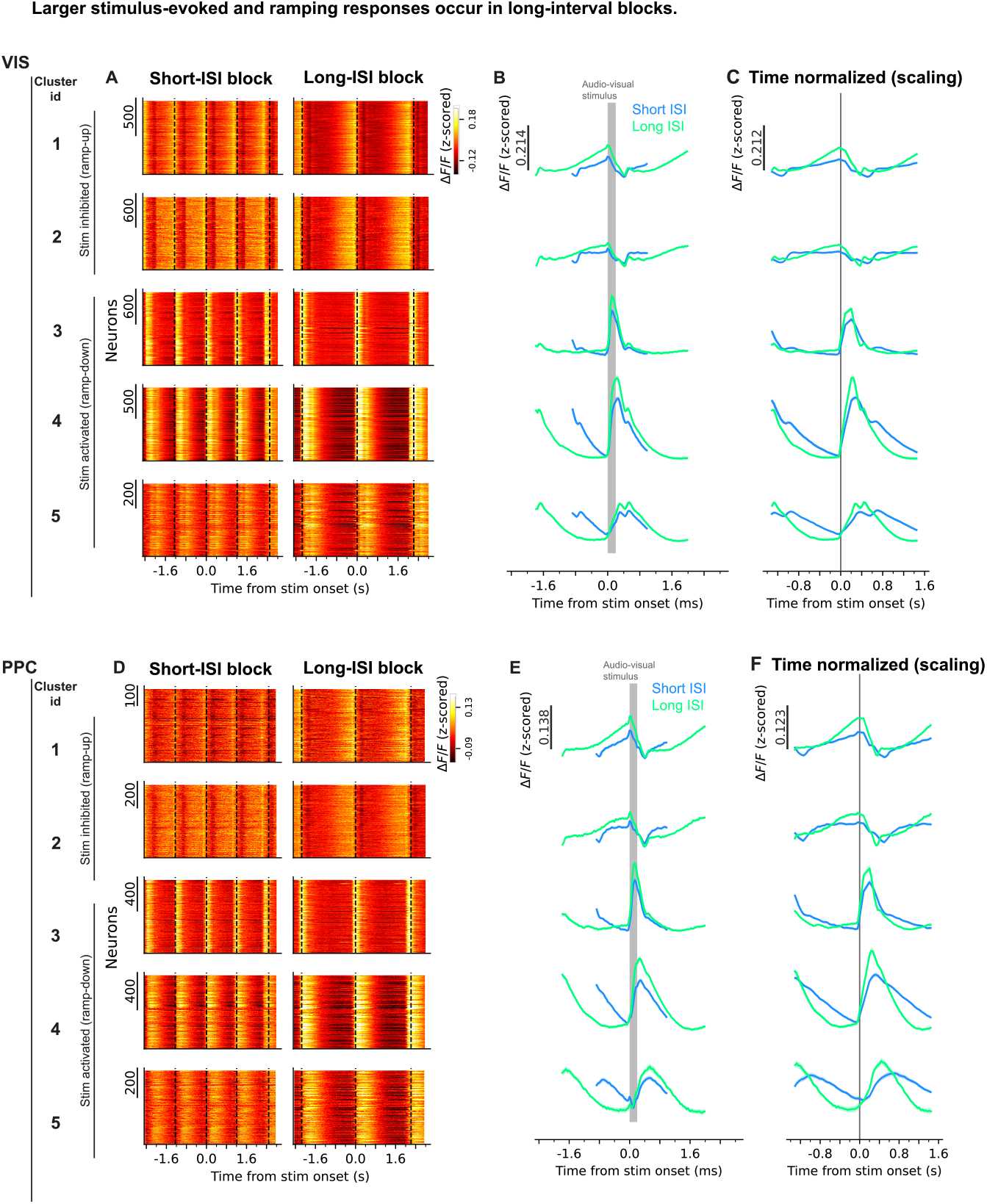
Larger stimulus-evoked and ramping responses emerge during longer interval blocks. Data from the short/long sessions, comparing responses after short vs. long intervals. Clusters are the same as in **Fig. 2**. Each row corresponds to a cluster. **VIS:** top panels; **PPC:** bottom panels. **(A)** Heatmaps of trial-averaged, single-neuron activity aligned on stimuli (vertical lines) in the short (**Left**) and long (**Right**) ISI blocks, for each cluster (top to bottom). Neurons are sorted identically for short and long blocks. **(B)** Neuron- and trial-averaged activity for short (blue) and long (green) blocks aligned on the stimulus (gray box). Mean ± 95% confidence interval, across pooled neurons and trials from all sessions of all mice. **(C)** Same as **(B)**, but with the time axis scaled to place short and long on the same timescale. **(D-F)** Same as **(A-C)**, but for PPC.

### Short/long interval transitions evoke immediate, relatively stable response changes

We next examined how responses changed following block transitions (**Fig. 4**). Across clusters, responses changed immediately after interval transitions. Following short-to-long transitions, stimulus-activated clusters increased response amplitude on the very first long-ISI trial (**Fig. 4**; S⟶L; VIS/PPC: rows 3-5), while stimulus-inhibited clusters exhibited much more pronounced ramps, already evident in the first long-ISI (**Fig. 4**; S⟶L; VIS/PPC: rows 1-2). The opposite effects were observed during long-to-short transitions, with stimulus-activated clusters decreasing in amplitude and stimulus-inhibited clusters showing attenuated ramps (**Fig. 4**; L⟶S).

**Figure 4.**
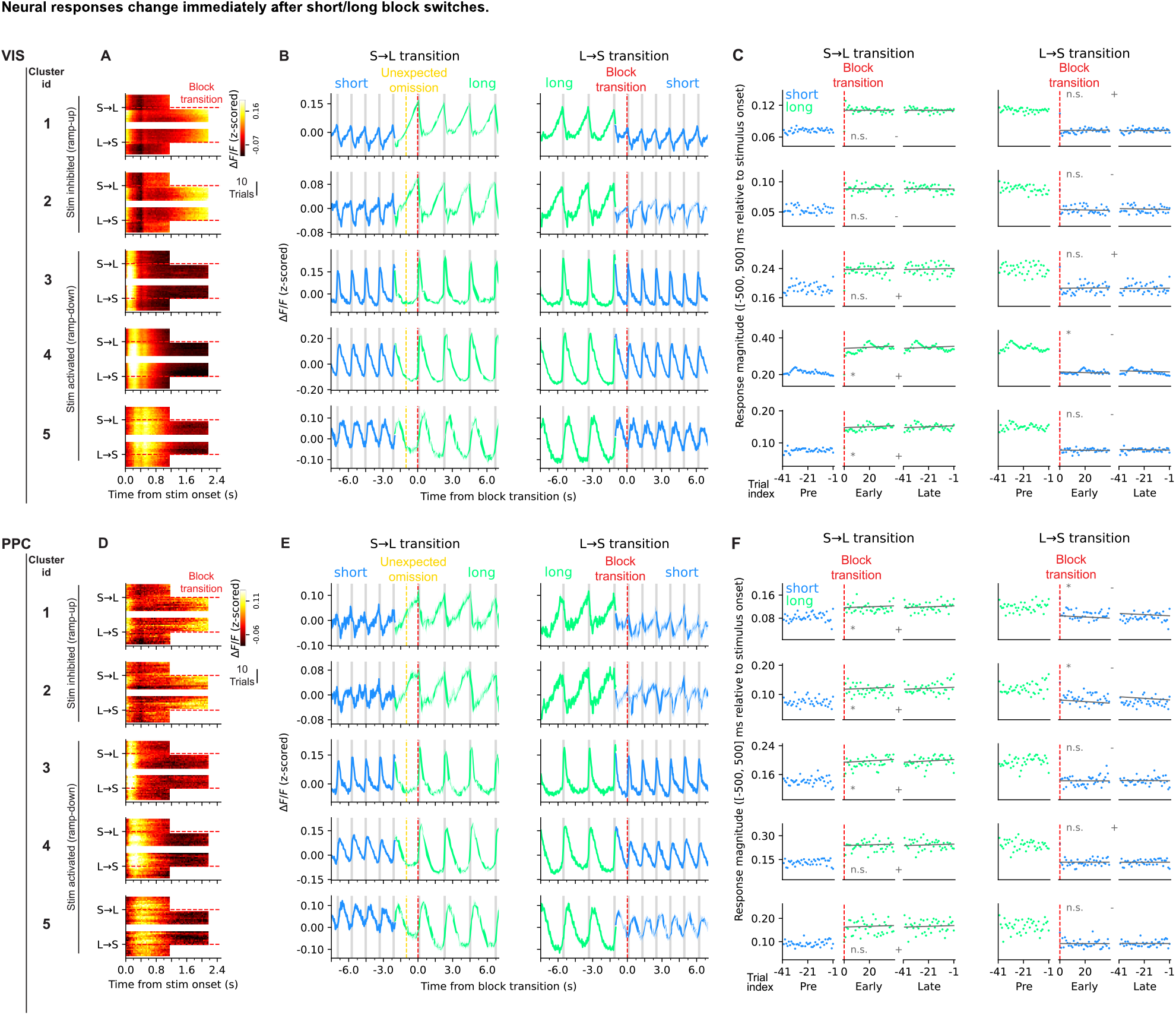
Interval transitions evoke immediate neural response changes that remain relatively stable throughout the block. Data from the short/long block sessions, comparing responses after short vs. long intervals. Clusters are the same as in **Fig. 2**. Each row corresponds to a cluster. **VIS:** top panels; **PPC:** bottom panels. **(A)** Heatmaps of trial-by-trial, population-averaged activity around block transitions (red horizontal lines), for both short-to-long (S→L) and long-to-short (L→S) block switches, for distinct clusters (top to bottom). Rows: individual trials. **(B)** Trial-by-trial neural activity around block transitions. Same as (**C)**, but shown as line plots (mean ± 95% confidence interval, across pooled neurons and block switches from all sessions of all mice). Red line: first stimulus after ISI switch for both transitions. Yellow line: expected stimulus timing based on the short block (first unexpected stimulus omission). Blue: short block; Green: long block. Gray boxes: stimuli. **(C)** Response-evoked magnitude was quantified in a ±500 ms window around stimulus onset as the difference between the upper and lower 30% of activity values. Magnitudes were computed for the 40 trials before the block transition (pre), immediately after transition (early), and at the end of the block (late). Linear fits were used to assess within-block changes over trials. Significantly positive or negative slopes are denoted by an asterisk, and +/− signs indicate slope direction. Each dot represents single-trial mean response, across pooled neurons and block switches from all sessions of all mice. (asterisk: p<0.05; n.s.: non-significant; two-sided t-test from *scipy*.*stats*.*linregress*). **(D-F)** Same as **(A-C)**, but for PPC.

To quantify these dynamics, we measured response magnitude across three epochs: immediately before a block transition, immediately after, and late within the block (Pre, Early, and Late, respectively; **Fig. 4C,F**). This analysis showed that the largest response change occurred immediately at the block transition. Response magnitudes remained relatively stable thereafter, although some clusters exhibited slight gradual within-block changes (**Fig. 4C,F**, fitted line, asterisk), consistent with mechanisms such as gradual adaptation buildup during repeated short-ISI stimulation and gradual recovery from adaptation during long-ISI blocks (see Discussion).

### Short/long interval transitions do not support temporal predictive processing

The predictive-processing hypothesis posits that a change in interval at block transitions should generate a temporal prediction error, which would gradually diminish across subsequent trials as new temporal predictions are learned. Our results contract this.

After short-to-long transitions, stimulus-inhibited clusters exhibited activity changes at the time of the unexpectedly omitted stimulus in the first long ISI (**Fig. 4B,E**; S⟶L; yellow line), which could superficially resemble a prediction-error signal. However, similar ramping activity persisted across later long-interval trials, rather than selectively decaying after the first unexpected omission (**Fig. 4C,F**; S⟶L).

Conversely, during long-to-short transitions, stimulus-inhibited neurons did not respond to the first short-ISI trial, and stimulus-activated neurons showed reduced response amplitude (**Fig. 4B,E**; L⟶S; red line).

Together, these results indicate that block transitions rapidly alter cortical response dynamics, but provide no evidence for transient temporal prediction-error signaling or gradual updating of temporal predictions.

### Fixed and jittered interval contexts evoke indistinguishable cortical dynamics

While our short/long-session results do not support temporal predictive processing, alternative explanations could still be considered. One possibility is that, following short-to-long transitions, prediction and prediction-error signals co-occur within the same neurons—prediction signals gradually increasing while prediction-error signals gradually decreasing—thereby canceling each other at the population level. Another possibility is that, following long-to-short transitions, an unusually high learning rate allows temporal predictions to update within a single trial, explaining the absence of gradual within-block increases in prediction-related activity.

To test these scenarios more directly, we performed fixed/jittered-ISI block sessions (**Fig. 1G**). Fixed blocks consisted of constant ISIs (1.5 s) with occasional unexpected short or long intervals (deviants: 0.5 s and 2.5 s). In jittered blocks, ISIs were drawn from a uniform 0.5–2.5 s distribution, so the same 0.5 s and 2.5 s intervals were expected. Predictive processing therefore predicts stronger temporal prediction-error responses to deviant ISIs in fixed compared to jittered blocks, as well as stronger temporal prediction signals in fixed blocks due to their greater temporal regularity.

Surprisingly, and in clear contrast to predictive-processing predictions, responses to deviant ISIs were indistinguishable between fixed and jittered blocks, for both short and long deviants (**Fig. 5**; VIS: **A,E;** PPC: **J,N**; Red: fixed; Blue: jittered; Arrow 2: unexpected stimulus omission; Arrow 3: unexpected stimulus presence). This key result provides strong evidence against temporal prediction-error coding in PPC or VIS during passive audio-visual stimulation. Likewise, pre-stimulus ramping activity in ramp-up neurons was nearly identical between fixed and jittered contexts (**Fig. 5A,E,J,N**; Arrow 1; see ramps prior to stimulus at t=0), inconsistent with ramping activity representing temporal prediction signals, which should have weakened under jitter.

**Figure 5.**
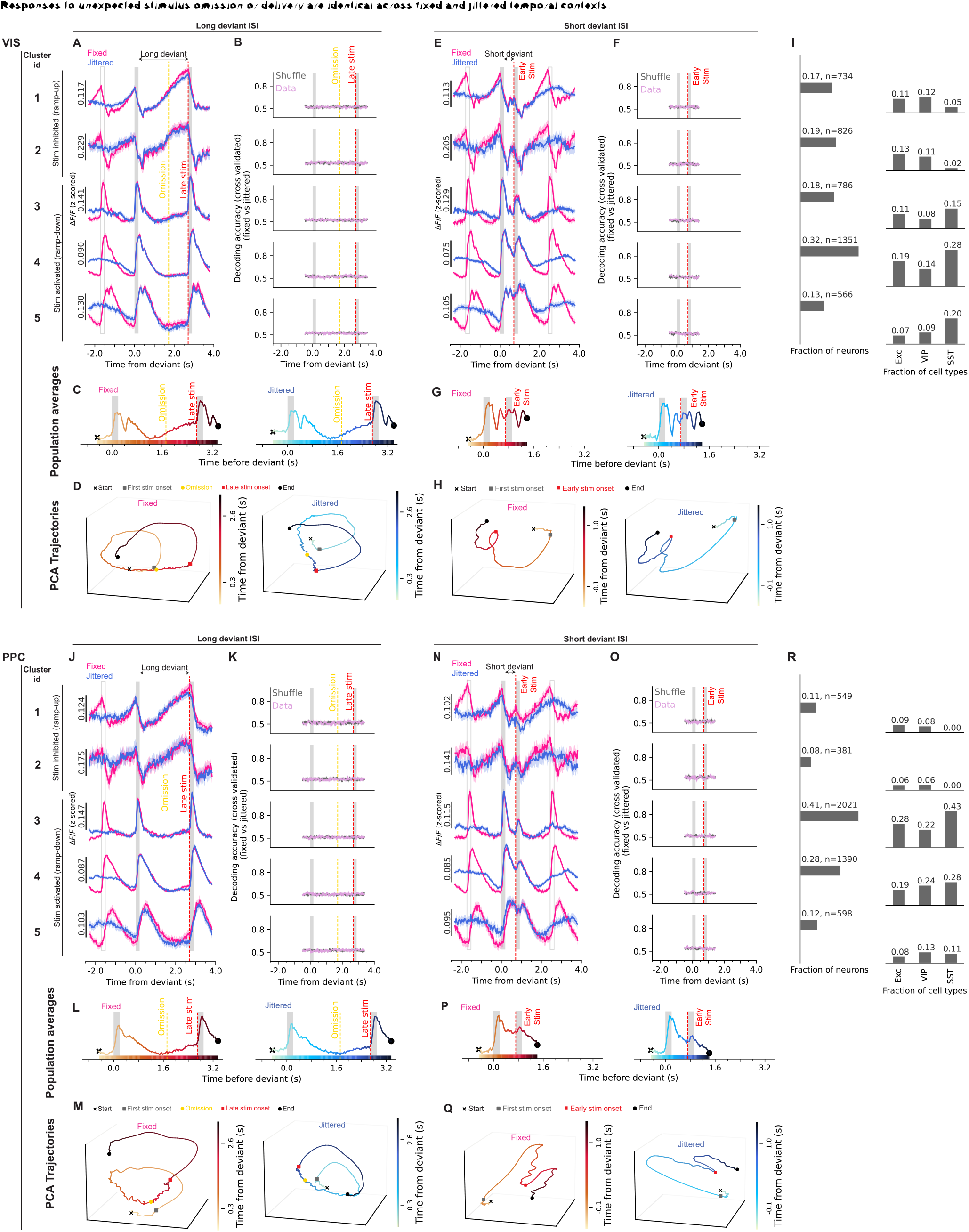
Neural responses do *not* differ between fixed and jittered ISI blocks for either long or short deviant ISIs. Data from fixed/jittered sessions, showing identical responses to both short and long deviant ISIs across fixed and jittered blocks. Clusters were identified on pooled neurons across all fixed/jittered sessions. **VIS:** top panels; **PPC:** bottom panels. Across fixed and jittered-ISI blocks, clusters 1–2 exhibited similar ramping activity preceding stimulus delivery (arrow 1) and following stimulus omissions (arrow 2). Clusters 3–5 exhibited similar stimulus-evoked responses following the unexpected stimulus delivery (arrow 3). Thus, neither pre-stimulus or omission-related ramping activity nor stimulus-evoked responses can be explained by temporal predictions or prediction-error signaling. **Fig. S4** quantifies single-trial responses to omissions and early/late stimuli, for each cluster. **(A)** Neural responses for each cluster, for fixed (red) and jittered (blue) blocks, aligned on the stimulus before the long deviant ISI. Mean ± 95% confidence interval, across pooled neurons and trials from all sessions of all mice. (Yellow line: expected but omitted stimulus; Gray boxes: stimuli; Empty boxes: stimuli in fixed and mean stimuli time in jittered). **(B)** Decoding accuracy for classifying fixed vs. jittered trials (cross-validated; Linear SVM; Mean ± 95% confidence interval, across cross-validated folds from all sessions of all mice; Purple: real data; Gray: shuffled data; Yellow line: omission). Separate decoders were trained at each time point using single-trial population activity, for each session. Decoder performance remained at chance across all time points, demonstrating similar responses between fixed and jittered conditions throughout the trial. **(C)** Population average, across all neurons of all clusters, for fixed (left) and jittered (right) blocks, aligned on the long deviant ISI. Traces are color-coded to match PCA trajectories in **(D)**. **(D)** PCA population trajectories for fixed (left) and jittered (right) trials, aligned on the long deviant ISI (projection on top 3 PCs); Yellow: time of the expected—but omitted— stimulus, based on the mean ISI at 1.5 sec; Red: unexpected late stimulus (stimulus after the long deviant ISI). PCA was performed on the combined fixed and jittered trial averaged population traces, across pooled neurons from all sessions of all mice. **(E–H)** Same as **(A–D)**, but aligned on the stimulus before the short deviant ISI. Red: unexpected early stimulus (stimulus after the short deviant ISI). **(I) Left:** fraction of all neurons assigned to each cluster. **Right:** fraction of neurons of each cell type assigned to each cluster, for fixed/jittered sessions. **(J-R)** Same as **(A-I)**, but for PPC.

To examine whether any prediction-error-like clusters were missed by our clustering approach, we revisited the hierarchical merging process used to define functional subtypes (**Fig. S5**). Across intermediate subclusters, we still did not observe any subpopulation that selectively responded to temporal deviations in fixed blocks compared with jittered blocks. Therefore, the absence of prediction-error responses is unlikely to reflect cluster extraction procedure washing out a selective prediction-error population.

To further test whether subtle differences might be obscured in population mean responses, we trained a linear decoder for each cluster to classify fixed versus jittered trials based on activity around deviant ISIs. In all clusters, decoder performance remained at chance, providing further evidence that responses to deviants were indistinguishable between fixed and jittered conditions (**Fig. 5B,K:** long deviants; **Fig. 5F,O:** short deviants; Purple: data; Gray: shuffled control).

A potential counter-argument is that both block types shared the same mean ISI (1.5 s), so average ramp dynamics might overlap despite differences in trial-to-trial variability. To directly examine this possibility, we computed a single-trial, neuron-averaged modulation index to quantify responses to unexpected events and compared its distribution between the two block types (**Fig. S4**; Methods). We found overlapping modulation-index distributions between fixed and jittered blocks for all deviant responses, including stimulus omissions during long deviants, and unexpected stimulus deliveries during short deviants or immediately following long deviants, as quantified by ROC analysis with area-under-the-curve (AUC) values consistently close to 0.5.

Prediction-error responses might be strongest during the earliest deviant presentations and therefore obscured by averaging across all deviant trials. To address this possibility, we performed an additional analysis using only the initial deviant trials (**Fig. S6**; Methods). We first selected strongly modulated neurons and then examined their average responses across only the first 10 deviant trials in fixed and jittered blocks. Even under this more sensitive analysis, we did not observe systematic differences in responses to deviant ISIs between fixed and jittered conditions (**Fig. S6**), indicating that even the earliest and strongest deviant-evoked responses were not detectably modulated by temporal predictability.

We also performed principal component analysis (PCA) to examine how population trajectories change during unexpected events in both fixed and jittered contexts. Both contexts exhibited similar trajectories: stimulus onset drove population to an evoked state, which then gradually relaxed back toward baseline (**Fig. 5**; VIS: **D,H**; PPC: **M,Q**; gray square: pre-deviant-ISI stimulus).

Importantly, in the fixed block, unexpected omissions did not perturb the trajectory, which continued in state space until the next stimulus, closely resembling the jittered trajectory (**Fig. 5D,M**; Yellow: omission; Red: late stimulus). Also, the late stimulus following the deviant ISI, despite its unexpected timing, produced a trajectory similar to that evoked by the predictable stimulus before the deviant ISI (**Fig. 5D,M**, gray vs. red squares), and this similarity was observed in both fixed and jittered conditions (**Fig. 5D,M**: PCA; **Fig. 5C,L**: population averages).

A similar lack of difference was observed for short deviants: in both fixed and jittered contexts, the post-deviant stimulus (“early stimulus,” **Fig. 5G,H,P,Q**; red) pushed trajectories into an evoked state. Compared to the stimulus before the short deviant (**Fig. 5G,H**; gray), this early stimulus produced a smaller trajectory change, consistent with the reduced response amplitudes in short versus long blocks (**Fig. 3B,E**).

Finally, we investigated whether cell-type interactions differ between fixed and jittered contexts. To address this, we leveraged our simultaneously imaged VIP– excitatory dataset and computed cross-trial Spearman pairwise correlations. Within and cross-population correlations evolved similarly in both fixed and jittered contexts (**Fig. S7**), suggesting that interactions between neurons are largely independent of temporal predictability in our passive paradigm.

Therefore, multiple complementary analyses—including precise cluster averaging, decoding, modulation-index quantification, correlation and principal component analysis—provided no evidence that temporal context influences responses to deviant intervals. Together, these findings argue against temporal predictive processing as the mechanism underlying pre-stimulus ramping activity or omission/deviant-related responses in VIS and PPC.

### Pre-stimulus ramping exists in naïve mice not exposed to regular intervals

To determine whether ramping responses depend on prior exposure to regular stimulus timing, we examined naïve mice that had never experienced regular stimulus intervals, and presented them with repeating audiovisual stimuli at random ISIs.

We observed the same neuronal clusters—stimulus-activated and inhibited—even in naïve mice (**Fig. 6**). Strikingly, pre-stimulus ramping activity was already present in the very first session (**Fig. 6C,K**; top row: ramp-up).

**Figure 6.**
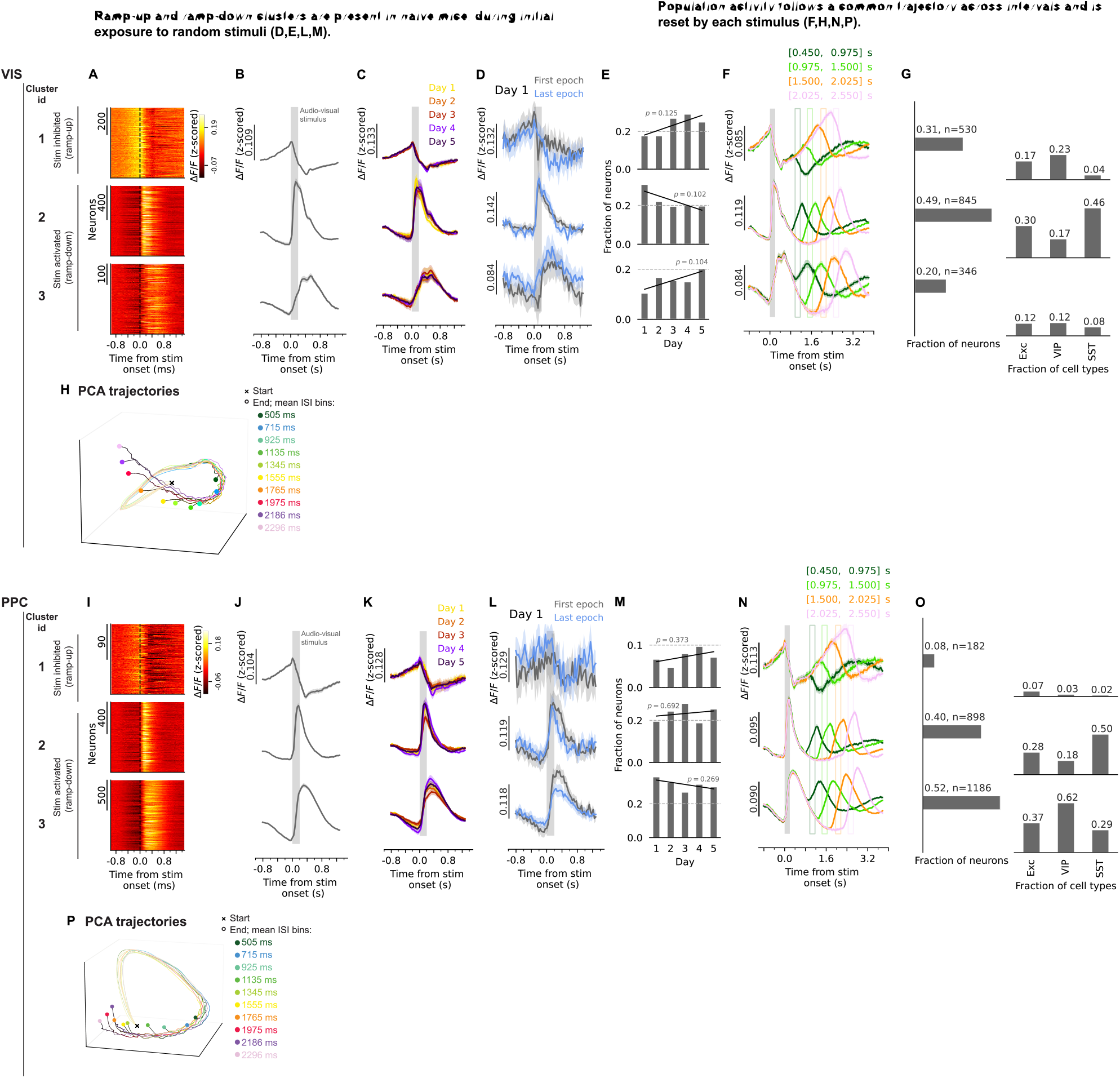
Neural ramps during repeating stimuli are intrinsic and reflect self-evolving sensory dynamics. Data from random sessions, showing that all neural clusters are present from the beginning in naïve mice, and do not extensive require learning to emerge. Clustering was performed on pooled neurons across all random sessions. Each row corresponds to a cluster. **VIS:** top panels; **PPC:** bottom panels. **(A)** Heatmaps of trial-averaged, single-neuron activity aligned on stimulus onset (black line), for each cluster (top to bottom). **(B)** Neuron- and trial-averaged responses aligned on the stimulus (gray box), for each cluster. Mean ± 95% confidence interval, across pooled neurons and trials from all sessions of all mice. **(C)** Averaged responses aligned on the stimulus, same as (B), but shown separately for individual random sessions (different colors). Mean ± 95% confidence interval. **(D)** Averaged responses for the first (gray) and last (blue) 50 trials of the first session. Mean ± 95% confidence interval, across pooled neurons and trials, from session 1 of all mice. See **Fig. S7A,C** for smaller trial-window averages. **(E)** Fraction of total neurons in each cluster distributed across the five random days. Linear fits were used to assess changes across days, with p values shown for each cluster. **(F)** Random ISIs divided into four bins (different colors). Mean ± 95% confidence interval, across pooled neurons and trials from all sessions of all mice. Neural responses averaged across trials within each ISI bin, showing how responses evolve across distinct ISIs. (Gray box: stimulus. Empty boxes: mean ISI of each bin). **Fig. S8B,D** groups trials based on the pre-stimulus ISI, showing how stimulus-evoked responses depend on the preceding interval duration. **Fig. S9** presents results from random sessions with a larger ISI range, showing that ramp-up responses plateau after ∼4sec. **(G) Left:** fraction of all neurons assigned to each cluster. **Right:** fraction of neurons of each cell type assigned to each cluster. **(H)** PCA of population activity (projection on top 3 components) from stimulus onset through the ISI. Random ISIs were divided into 10 bins (210 ms each), and trajectories are shown up to the mean ISI for each bin. Trajectories follow a common path across intervals and extend further along it with longer ISIs. PCA was done on trial-averaged population responses, for each ISI bin. **(I-P)** Same as **(A-H)**, but for PPC.

To address the possibility that rapid predictions might emerge within the first day, we compared neural responses between the first 50 and last 50 stimuli of day 1, but again observed similar ramping responses (**Fig. 6D,L, Fig. S8**; black: first epoch; blue: last epoch). The prevalence of clusters remained generally stable across days (**Fig. 6E,M**).

We further examined smaller trial windows to better estimate when pre-stimulus ramping first emerged (**Fig. S8A,C**). In VIS, ramping activity was already evident within the first 20 stimuli on day 1, whereas 10-trial windows were too noisy to reliably resolve early versus late epoch differences (**Fig. S8A**; top row). In PPC, pre-stimulus ramping appeared to be present within the first 30 trials, but the traces were noisier than in VIS— likely due to the lower prevalence of stimulus-inhibited neurons (**Table 2,3**)—thereby, preventing clear conclusions (**Fig. S8C**; top row).

Overall, these results indicate that ramping activity does not require extensive within-session learning, as it was already apparent within ∼20 trials in VIS and ∼30 trials in PPC. However, rapid emergence during the very first few trials cannot be excluded (see Discussion).

Therefore, multiple complementary analyses—including precise cluster averaging, decoding, modulation-index quantification, and PCA—provided no evidence that temporal context influences responses to deviant intervals. Together, these findings argue against temporal predictive processing as the mechanism underlying pre-stimulus ramping activity or omission-related responses in VIS and PPC.

### Ramping activity arises from intrinsic sensory dynamics not learned predictions

Across all paradigms, our evidence argues that neural responses—and particularly the ramp-ups—do not reflect temporal prediction signals. A key question, then, is what underlies the response changes during short/long interval block switches (**Fig. 4**), or following deviant ISIs (**Fig. 5**). To address this, we binned the random intervals and examined how responses varied across distinct interval lengths (**Fig. 6F,N**).

Stimulus-activated responses were similar across interval bins, but their relaxation (ramp-down) was truncated when the next stimulus occurred early (**Fig. 6F,N;** rows 2-3; green shades) and extended further when the next stimulus was delayed (**Fig. 6F,N**; rows 2-3; orange/pink). Likewise, the ramp-up in stimulus-inhibited clusters was interrupted if a stimulus occurred early, whereas it continued further when the next stimulus was delayed (**Fig. 6F,N**; row 1; Green shades: shorter bins; Orange/pink: longer bins). Thus, neural responses across ISI bins appeared as different snapshots of the same underlying dynamic, interrupted and reset whenever the next stimulus arrived (**Fig. 6F,N**).

Population trajectory analysis confirmed this view: trajectories followed a common path across all intervals, simply evolving further along the same trajectory until reset by the next stimulus (**Fig. 6H,P**; random ISIs divided into 10 bins; Trajectories start at stimulus onset and end at the mean ISI bin).

To examine how long ramping activity can persist, we extended the random-ISI paradigm to longer intervals (3-7.5 s; **Fig. S9**). Ramp-up activity increased monotonically for the first ∼4 s after stimulus onset before reaching an apparent plateau.

Together, these results indicate that VIS and PPC ramping dynamics preceding stimuli or following unexpected stimulus omissions (long deviant ISIs; **Fig. 4,5**: rows 1– 2) reflect intrinsic relaxation from the stimulus-evoked state, rather than temporal prediction or prediction-error mechanisms.

### Heterogeneous rise/decay kinetics enable robust population coding of time

Our results so far demonstrate that post-stimulus activity reflects the unfolding of intrinsic cortical dynamics (**Fig. 6**), independent of predictive processing (**Fig. 4,5**). We next asked what information these ramps might carry—specifically, whether they encode elapsed time (**Fig. 1A**). We reasoned that the diversity of rise and decay kinetics across neurons (**Fig. 2**) should support a robust population code for time: neurons with different temporal profiles would peak at distinct moments, collectively spanning a wide range of intervals.

To assess this, we trained decoders to discriminate pairs of time bins based on population activity within each cluster (**Fig. 7B,H**; cross-validated linear SVM trained on long-block data from the short/long sessions). PCA was performed on each cluster’s population activity to reduce the population dimension to 50. PCA was first applied to each cluster’s population activity to reduce dimensionality to 50 components while preserving >99% cumulative explained variance (**Fig. 2F,L**). Using the same dimensionality across clusters enabled comparison of decoder performance between populations containing different numbers of neurons (Methods).

**Figure 7.**
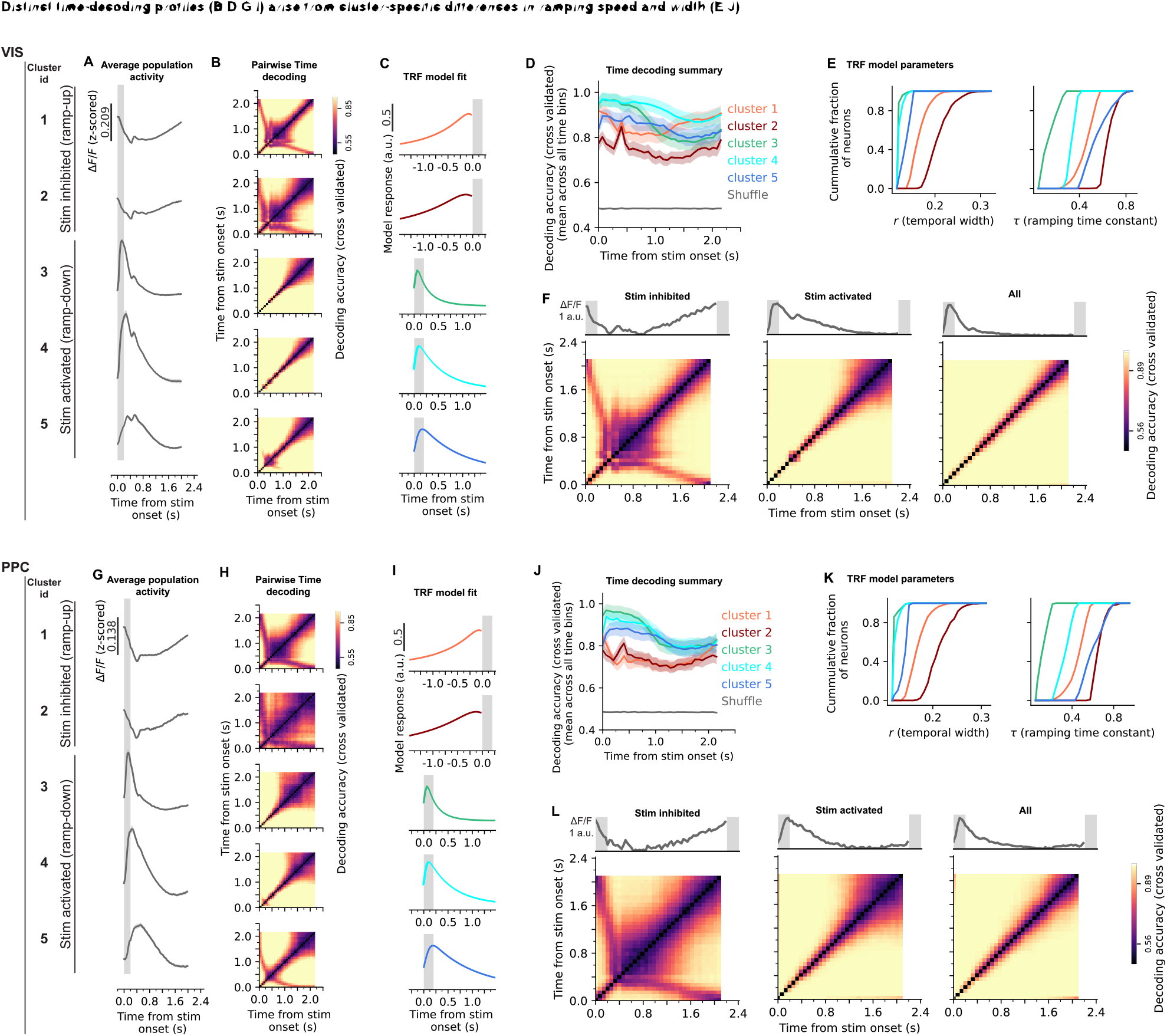
Heterogeneous ramp kinetics enable robust population coding of time. Data from the long block of short/long sessions, showing heterogeneous kinetics across clusters, which leads to distinct temporal coding profiles across the interval. Clusters are the same as in **Fig. 2**. Each row corresponds to a cluster. **VIS:** top panels; **PPC:** bottom panels. **(A)** Mean population activity, shown for each cluster (top to bottom). **(B)** Pairwise classification accuracy matrices (cross-validated). Each cell shows the accuracy for distinguishing a given pair of time bins, shown for each cluster. Mean across cross-validated folds. For decoding analyses, PCA was first applied to each cluster’s population activity, followed by a linear SVM trained on the top 50 principal components. See Methods for details. **(C)** Temporal receptive field model response (see **Fig. 1I**), averaged across neurons, shown for each cluster. Orange, red: ramp-up clusters. Green, cyan, blue: ramp-down clusters. **(D)** Decoding accuracy at each time point, computed by averaging the pairwise classification accuracies in panel B across all other time bins (excluding the diagonal). This quantifies how accurately each time point can, on average, be discriminated from the rest of the trial. Mean ± SEM across cross-validated folds. Orange, red: ramp-up clusters. Green, blue, cyan: ramp-down clusters. **(E)** Distribution of model parameters across neurons within each cluster; Left: temporal width (r); Right: ramping time constant (τ). Distinct decoding profiles **(B,D)** arise from cluster-specific differences in ramping dynamics. **(F)** Pairwise temporal classification accuracy matrices (cross-validated) for all stimulus-inhibited neurons (left), all stimulus-activated neurons (middle), and the combined population containing both response classes (right). **Top:** Population average for corresponding neurons. **Bottom:** Pairwise classification accuracy matrices, similar to (B). Temporal decoding performance is enhanced in the heterogeneous population containing both ramp-up and ramp-down neurons. **(G-L)** Same as **(A-F)**, but for PPC.

All clusters carried information about elapsed time—as expected from any time-varying signal—but their decoding profiles differed systematically (**Fig. 7;** VIS: **B,D**; PPC: **G,I**; chance at 50%). Stimulus-activated clusters decoded short intervals more accurately, with high initial accuracy that declined over time (**Fig. 7D,J**: green, cyan, blue). In contrast, stimulus-inhibited clusters showed stable decoding throughout the interval, with a small increase in accuracy near the end, consistent with their ramp-up dynamics (**Fig. 7D,J**: orange, red).

To understand how response kinematics shape these profiles, we compared parameters of the temporal-receptive-field model across clusters. A clear pattern emerged: clusters with faster and narrower responses supported higher temporal precision (**Fig. 7E,K**; green; cyan), whereas clusters with slower and broader responses provided more stable decoding (**Fig. 7E,K**; orange; red).

Finally, we assessed whether heterogeneous response kinetics improve temporal decoding at the population level. We first pooled all stimulus-inhibited clusters and performed decoding analyses (**Fig. 7F,L**; left), then repeated the analysis for pooled stimulus-activated clusters (**Fig. 7F,L**; middle), and finally combined neurons from both response classes (**Fig. 7F,L**; right). Temporal decoding performance was substantially enhanced in the heterogeneous population containing both response classes (**Fig. 7F,L**; right). Importantly, PCA dimensionality reduction was applied prior to decoding to ensure that this improvement did not simply reflect the larger population size of the combined dataset. These findings demonstrate that populations containing heterogeneous temporal dynamics enable more accurate decoding of elapsed time.

Altogether, our findings support a simple mechanistic model: stimulus onset resets the population into an evoked state that then relaxes toward baseline. Both the rise into the evoked state and the decay back vary across neurons. This diversity in response kinetics enables the population to represent elapsed time across multiple scales. Accordingly, <u>changes in neural responses across ISIs—including unexpected stimulus omissions—reflect sampling different phases of a shared trajectory, rather than adaptive or predictive mechanisms.</u>

## DISCUSSION

Our central finding is that the ramping activity in visual and parietal cortices—observed during passive repeating audiovisual stimulation—emerges from the ongoing evolution of stimulus-evoked dynamics, rather than from temporal predictive processing. We further demonstrate that these ramping responses have heterogeneous kinetics, enabling an intrinsic, distributed representation of elapsed time. Whether mice actually read out this temporal information remains unknown, as our paradigm was passive and did not require a behavioral report. Nevertheless, the fact that elapsed time can be decoded from these responses suggests that downstream circuits could access this signal to estimate time even in the absence of explicit task demands to attend to time.

More specifically, our analyses revealed that during passive presentation of repeated audiovisual stimuli in awake mice, neurons in the medial visual cortex (VIS) and posterior parietal cortex (PPC) segregated into two main functional categories: stimulus-activated (ramp-down) and stimulus-inhibited (ramp-up) neurons. Within each category, subtypes were distinguished by their ramp kinetics, spanning fast to slow temporal profiles **(Fig. 2)**. Three findings showed that these ramps were independent of temporal prediction or prediction-error processes: **(1)** neural activity changed immediately across blocks with short and long ISIs, and remained relatively stable thereafter (**Fig. 4**); **(2)** responses immediately before expected stimuli and after unexpected onset shifts were identical in fixed and jittered blocks (**Fig. 5**); and **(3)** the same functional clusters were present in naïve mice during fully random ISIs (**Fig. 6**).

Analysis of responses as a function of interval length (**Fig. 6F,H**) showed that apparent differences across short and long intervals simply reflect the intrinsic evolution of stimulus-evoked activity, rather than responses to omitted or predicted events. Decoding analyses further revealed that these clusters collectively encode elapsed-time information, with each cluster’s interval preference determined by its characteristic ramp kinetics (**Fig. 7**).

### Visual omission responses: comparison with prior studies

Our findings demonstrate that ramping activity during unexpected omission of visual stimuli, in visual and parietal cortex, does not reflect temporal prediction-error signals (*41-43, 49*), but instead reflects the intrinsic evolution of stimulus-evoked activity as it relaxes between successive stimuli. Moreover, although behavioral variables may co-vary with omissions (*43*), our results show that they are not the cause of these responses— fundamentally, during passive viewing conditions, there is no discrete “omission response” in visual and parietal cortex, only the continuation of ongoing stimulus-evoked dynamics.

A key distinction between this and prior studies lies in how *temporal predictive processing* and *omissions* are defined. Here, temporal prediction refers specifically to the brain’s estimate of *when* the next stimulus will occur. We manipulated this by jittering the duration of a neutral gray interval between otherwise identical stimuli—holding stimulus identity and duration constant—to isolate timing-related processes without confounding changes in visual content or duration.

In contrast, image-sequence-based studies (*50, 67*) have defined an “omission” as repeating the previous image instead of presenting the next. Consequently, prediction errors in these paradigms reflect violations of both stimulus identity and duration within a learned image sequence. Robust “omission” effects in those studies emerge only after multi-day training—specifically, exposure to the sequence without deviants following an initial reference session (*50*)—indicating sequence-specific learning.

Our paradigm, by contrast, employs simple repeated-image presentations and defines omission as a prolonged neutral-gray period, thus probing *timing per se* without changes in stimulus identity or duration. Importantly, even after repeated daily exposure to sessions containing rare omissions, we did not observe the emergence of omission-related activity, suggesting that these dynamics do not depend on learning the session’s temporal structure. While we cannot entirely rule out that omission responses might emerge if animals were first trained on sessions without omissions, we consider this very unlikely given the quite low omission rate in our paradigm and the absence of a complex image sequence that could support such learning.

Other studies have examined omissions in closed-loop visuomotor contexts, where visual flow is coupled to locomotion (*44, 46, 68*). In those tasks, omission events—replacing an expected image with gray—represent violations of *sensorimotor contingencies* rather than purely temporal predictions. Thus, while the term “omission” is shared, the underlying predictive computations differ fundamentally.

A recent study most comparable to ours (*49*), reanalyzed large-scale Allen Institute datasets and concluded that omission responses in visual cortex do not represent prediction errors, but may instead reflect temporal expectation or rebound activity. Our results extend these insights using dedicated experimental paradigms that directly dissociates elapsed-time encoding from prediction. By systematically varying interval regularity and experience, we confirm that visual cortical activity—during passive behavior—neither signals temporal prediction errors nor predictions themselves. Rather, ramping dynamics reflect intrinsic, time-evolving processes within local circuits.

Taken together, these observations emphasize the need for precise definitions of “omission” and “temporal predictive processing” when designing experiments and interpreting omission-related activity across sensory and behavioral contexts.

### Possible sites of temporal predictive processing during visual perception

A pressing question raised by our findings is where in the brain temporal predictive processing during passive visual inputs is implemented (*3*). Hippocampal neurons—and subcortical regions such as the thalamic medial geniculate nucleus—have been suggested to carry temporal prediction-error signals, since, compared to the gradual ramps observed in visual cortex, they exhibit steeper responses to visual omissions (*49*).

Another potential candidate is the cerebellum. Numerous studies have established its role in encoding timing and error signals during sensorimotor behavior (*18, 22, 23, 69-75, 76 Li 2018*). Fewer studies have examined the cerebellum’s role during passive stimulation (*77*), but evidence shows that the cerebellum is active following unexpected omissions of passive somatosensory stimuli (*78*) and is necessary for detecting temporal mismatches during passive auditory inputs (*79*). With respect to visual inputs, the cerebellum encodes repeated visual stimuli and their omissions during motor behavior (*80*), but its role during passive visual perception remains to be determined.

Other candidate regions for temporal predictive processing include the basal ganglia (*19, 20, 80*) and the prefrontal cortex (*9, 30, 31*), alongside the auditory cortex, whose role in auditory temporal predictive processing is well established (*33, 81, 82*).

VIS/PPC, too, may reflect temporal prediction signals but in active perceptual paradigms where animals need to attend to stimuli to receive reward. Human EEG studies support this view: anticipatory signals were absent during passive rhythmic streams but emerged when subjects prepared for action (*83*). Future studies using paradigms similar to ours, but including active perceptual or behavioral contexts, will be critical for determining whether VIS and PPC dynamics shift from elapsed-time coding toward predictive processing. Also, several animal studies have reported predictive and error signals in cortical areas when sensory events are behaviorally relevant (*84-89*). This implies that task engagement may be necessary to recruit temporal predictive processing in these circuits.

### The importance of incorporating multiple experimental paradigms when interpreting predictive processing results

A common approach for studying temporal predictive processing is to present stimuli at regular intervals. However, in such paradigms, temporal prediction signals often overlap with elapsed-time signals, which reflect the time since the previous stimulus. This overlap makes it difficult to determine whether neural activity reflects prediction or simple temporal tracking.

Our fixed/jittered paradigm with deviant ISIs—together with the random-ISI paradigm in naïve mice and alternating short/long block paradigm—provide a powerful tool to separate elapsed-time signaling from temporal predictive processing. <u>Without these paradigms, neural responses to long-ISI deviants (omissions) could be mistakenly interpreted as temporal prediction errors, and ramping signals as temporal predictions.</u>

Our results demonstrate that ramping and omission responses—in visual and parietal cortices during passive stimulation—are not temporal prediction signals but the intrinsic evolution of stimulus-evoked activity (**Fig. 6**) that carries linearly decodable information about elapsed time (**Fig. 7**). Similar “population clock” dynamics have been described across brain regions (*31, 34, 90*), indicating that temporal coding can arise intrinsically from network dynamics without prediction or reinforcement.

### Ramp-up dynamics may undergo slower experience-dependent changes across days

Although pre-stimulus ramping was already evident within the first ∼20 stimulus repetitions in naïve mice (**Fig. S8**), we cannot exclude the possibility that these dynamics are absent during the very first few trials and emerge rapidly thereafter.

Consistent with this possibility, the fraction of excitatory and VIP neurons exhibiting ramp-up dynamics gradually increased across paradigms, while the fraction of stimulus-activated neurons decreased. This effect was especially pronounced in PPC, but was also evident in VIS (**Table 2,3**). In VIS, the fraction of stimulus-inhibited neurons across paradigms—in order of presentation to mice—was 31%, 36%, 44%, and 64% for the random, fixed/jittered, short/long, and extended-ISI random paradigms, respectively. In PPC, these fractions were 8%, 19%, and 30% for the random, fixed/jittered, and short/long paradigms, respectively (random**: Fig. 6G,O**; **Fig. 5I,R**: fixed/jittered; **Fig. 2C,I**: short/long; **Fig. S9F**: extended-ISI random).

While our fixed/jittered and short/long paradigms argue against these changes reflecting learning of temporal predictions, the gradual increase in ramp-up neurons may instead reflect slower experience-dependent circuit changes, such as alterations in sensory gain or neuromodulatory inputs across repeated passive exposure over days. Consistent with this interpretation, our previous study showed that VIP neurons in primary and higher visual cortices transition from stimulus-activated to stimulus-inhibited responses as initially novel images become familiar (*42*).

Additionally, omission-related VIP/SST dynamics in visual cortex are strongly modulated by behavioral strategy (*91*), supporting the view that cortical ramping dynamics are shaped by experience and internal state. Future experiments directly monitoring behavioral state and arousal longitudinally across days may help clarify the underlying mechanisms.

### Ramping activity emerges from stimulus-reset attractor dynamics

Our results can be understood within an attractor-dynamics framework (*92-94*), in which each stimulus pushes neurons from baseline into a transient evoked state, after which activity relaxes back toward baseline. In this view, responses differ across interval lengths not because of trial-by-trial learning, but because the same intrinsic trajectory is reset and restarted at different points in its cycle.

This framework naturally accounts for our key findings: **(1)** the immediate changes in neural responses at block transitions (**Fig. 4**) without invoking trial-by-trial learning; **(2)** identical responses between fixed and jittered ISI contexts (**Fig. 5**), despite lower temporal predictability in the jittered case; and **(3)** the presence of the same neural clusters in both naïve and experienced mice (**Fig. 6**).

Importantly, this view situates our results within a broader attractor literature. Stimulus-reset attractor dynamics have been linked to variability quenching at onset (*93, 95*) and to persistent activity in working-memory and decision circuits (*96-98*). Our findings extend this framework to interval signaling: stimulus onsets place VIS/PPC populations onto reproducible evoked trajectories whose cluster-specific kinetics tile elapsed time, thereby yielding a learning-independent population code for interval duration (*8, 10*).

### Diverse neuronal dynamics provide a robust, distributed code for time

Neurons in VIS and PPC showed heterogeneous dynamics after stimulus presentation: both the rise into the evoked state and the decay back to baseline varied across cells. This heterogeneity aligns with reservoir models of diverse timescales, where variable neuronal kinetics create overlapping trajectories that together form a distributed population code for elapsed time (*8-10, 15, 16*).

Evidence for such diverse (*58*) timescales extends across brain systems. In the cerebellum, granule cells provide heterogeneous temporal kernels important for interval learning (*18, 23, 99*); in the striatum, medium spiny neurons exhibit sequential dynamics that tile time over seconds (*19, 20*); and in the hippocampus, “time cells” fire at distinct delays from an event, collectively spanning behavioral intervals (*17, 21*).

The diversity of rise and decay dynamics effectively provides a set of temporal filters in which each neuron is tuned to a different portion of the interval. In this sense, the population acts like a basis set: overlapping response profiles that can be flexibly combined downstream to reconstruct elapsed time.

Such an organization enhances robustness to noise, since time is represented redundantly, and allows different circuits to emphasize narrow or broad signals depending on task demands. Consistent with this idea, combining ramp-up and ramp-down populations substantially improved temporal decoding performance compared to either population alone (**Fig. 7F,L**), demonstrating that heterogeneous response kinetics enhance the robustness and accuracy of distributed temporal representations.

<u>This framework parallels basis-function models of time in recurrent networks and supports the view that elapsed time is encoded through a distributed population code rather than by a single dedicated mechanism</u> (*10, 13, 15*).

### Cell-type-specific mechanisms underlying heterogeneous neural dynamics

While all cell types contributed to each functional cluster, there were notable cell-type-specific biases. SST neurons were predominantly stimulus-activated, whereas excitatory and particularly VIP neurons were more prevalent in the stimulus-inhibited cluster (**Fig. 2D,J**). Given that our stimuli were presented at high contrast, these results are consistent with prior findings (*100*) showing that SST neurons preferentially respond to high contrast, whereas VIP interneurons and layer 2/3 excitatory neurons are more responsive to low contrast.

The stimulus-inhibited (ramp-up) neurons observed here could reflect bottom-up inputs from circuits containing *suppressed-by-contrast* cells (*101*) previously described in the retina (*102*) and thalamus LGN (*103*), or alternatively, modulation by top-down influences such as behavioral state, reinforcement, or neuromodulatory drive (*104-108*).

The heterogeneous rise and decay dynamics we observe can be explained by a combination of intrinsic, synaptic, circuit, and modulatory mechanisms, with E/I (excitatory/inhibitory) motifs playing a key role in shaping this diversity, as outlined below.

At the intrinsic level, neurons differ in their ion channel composition (*109, 110*). Synaptic inputs also vary in their filtering properties: differences in release probability, vesicle pool size, and receptor kinetics produce characteristic time constants, with some synapses showing rapid depression and others facilitation, and receptor subtypes contributing fast or slow components (*111, 112*).

Beyond single-cell properties, the E/I balance in recurrent circuits amplifies this diversity, as neurons within inhibitory-dominated loops recover quickly, while those in excitatory-rich loops maintain activity longer.

Finally, neuromodulators further shape kinetics in a cell-type-specific manner: acetylcholine recruits VIP interneurons to gate inhibition in VIS (*106, 107*), noradrenaline broadens integration times and enhances ramping dynamics (*113*), and dopamine adjusts recurrent integration in prefrontal networks (*114*).

### Recovery processes may underlie interval-dependent response scaling

The larger response amplitudes observed during long-ISI blocks, across excitatory, SST, and VIP populations (**Fig. 3B,E, Fig. S9B,D**), likely arise from recovery processes operating on sub-second to multi-second timescales. Longer intervals allow partial restoration of synaptic and circuit resources depleted by repeated stimulation.

Mechanistically, this could reflect **(1)** replenishment of vesicle pools following synaptic depression (*115*), **(2)** relaxation of activity-dependent gain control that suppresses network responses at short ISIs (*116*), **(3)** recovery from intrinsic spike-frequency adaptation that limits excitability at short intervals (*117*), and **(4)** recovery of PV/SST synaptic transmission, leading to stronger transient inhibition and consequently larger rebound responses in excitatory neurons at longer intervals (*118*). Together, these slow restorative dynamics can account for the interval-dependent amplitude scaling without invoking predictive or learning-based mechanisms.

### Mixed-response neurons reveal a continuum of cortical ramping dynamics

A subset of neurons (“mixed” neurons, ∼37%) was not strictly classified as ramp-up or ramp-down by the TRF model and was therefore excluded from the primary figures (**Fig. S3**). Importantly, these neurons led to the same overall conclusions as the main populations; notably, their responses remained similar across fixed and jittered conditions (**Fig. S3;** VIS: **G,H;** PPC: **P,Q**).

Inspection of their average response profiles (**Fig. S3D,M**) suggests that these mixed neurons exhibited multi-component responses following each stimulus: an initial brief stimulus-evoked response followed by inhibition, then a delayed secondary response (∼0.4 s after stimulus onset), and finally gradual ramping dynamics. Thus, in these neurons, ramping activity may not simply reflect relaxation from stimulus inhibition. In addition, their overall response amplitudes were substantially smaller than those of the main classified clusters, which may have contributed to less reliable TRF classification.

Together, these observations suggest that cortical ramping dynamics likely exist along a continuum of temporal response motifs rather than as two perfectly discrete response classes. Importantly, however, even these mixed-response neurons did not exhibit signatures consistent with temporal predictive processing.

### Limitations of the study

1. We used two-photon calcium imaging, which provides only an indirect proxy of spiking activity. The slow kinetics of calcium indicators blur fast transients, potentially biasing the estimation of timescales and attenuating high-frequency components of neural responses (*57, 119*). These factors may influence the precise shape and timescale of the ramp-like signals reported here. Nonetheless, qualitatively similar slow ramp-like dynamics during repeated visual stimulation have also been observed in electrophysiological recordings from visual cortex (*49, 120*), suggesting that such dynamics are not specific to calcium-imaging measurements.
2. Viral injections were targeted to medial visual cortex using stereotaxic coordinates. Post-hoc histology confirmed GCaMP expression in the medial part of the visual cortex (area PM; **Fig. S1**, top). PM is considered a higher visual area with a preference for higher spatial frequencies and lower temporal frequencies (*121, 122*). Although response properties can differ across visual areas, our initial experiments (unpublished data) targeting the primary visual cortex (V1) yielded similar results to those observed in the medial visual cortex in this study. Taken together with the lack of difference between the medial visual cortex and PPC—an associative area linking stimulus and action—we therefore consider it unlikely that our core finding (stimulus-reset ramps invariant to temporal predictability) depends on the precise visual subarea. Yet, additional experiments directly targeting V1 could further clarify its role in temporal predictive processing.
3. While stimulus orientation was varied across tens of trials, the present study focused specifically on temporal structure rather than stimulus identity. Our findings therefore do not address prediction or prediction-error signals related to *what* stimulus appears, but only those related to *when* it appears.
4. We restricted our analyses to layer 2/3 neurons in VIS and PPC; therefore, our data does not rule out predictive computations in other laminae. Predictive signals may be distributed across cortical layers or expressed preferentially in deeper-layer output circuits, including layer 5 neurons with distinct cortico-cortical, cortico-striatal, cortico-thalamic, and subcortical projection patterns (*123-125*).
5. Our cell-type coverage was also limited to excitatory, SST, and VIP neurons. Other inhibitory populations such as NDNF interneurons—shown to disinhibit PV cells and gate long-range inputs in layer 1—likely shape predictive signals and could play a role in temporal predictive processing (*126, 127*).

### Discussion summary

Neural dynamics in VIS/PPC during passive audio-visual stimulation do not support temporal predictive processing. Instead, they are consistent with intrinsic stimulus-reset relaxation dynamics whose diverse timescales enable distributed temporal signaling. Other brain regions, such as the hippocampus and the cerebellum, remain potential candidates for temporal predictive processing during passive behavior. Finally, prior work suggests that active engagement, involving top-down inputs and reward contingencies, can recruit prediction and error signals in sensory areas, raising the testable hypothesis that task demands may shift visual/parietal responses from intrinsic elapsed-time coding toward explicit temporal predictive processing.

## MATERIALS AND METHODS

### Use of AI-assisted coding tools

Portions of analysis code were developed and refined with assistance from OpenAI GPT-5.1 and later versions. All AI-assisted code was reviewed, tested, modified, and validated by Y.H. prior to use. F.N. carefully reviewed and evaluated all analyses and resulting outputs.

### Subjects

Female and male transgenic mice were used in the current study. Mice included VIP-tdTomato and SST-cre (Jackson 013044) lines. VIP-tdTomato line was produced by breeding VIP-cre (Jackson 031628) with Ai14 (Jackson 007914; cre-dependent tdTomato). All procedures were approved by the Institutional Animal Care and Use Committee at the Georgia Institute of Technology and agreed with guidelines established by the National Institutes of Health.

### Sample size and statistical tests

Ten mice (female and male) were used: 5 for PPC imaging and 5 for VIS imaging. 18,904 VIS neurons and 14,835 PPC neurons were imaged across all sessions (**Table 1**). Imaged neurons included genetic subtypes: SST, VIP, and non-VIP (presumptive excitatory). Each genetic subtype was further divided into functional classes (stim-inhibited, stim-activated, mixed; **Table 2,3**). The mixed class was excluded from the main figures but quantified in **Fig. S3**.

Statistical tests are specified in the corresponding Methods sections and/or figure captions. Statistical significance was defined as p < 0.05.

### Behavioral paradigms

Head-fixed mice were seated in a half-cylindrical tube (*128*), and received 3 session types (behavioral paradigms): random, fixed/jittered, and short/long sessions. Each mouse received 5 random sessions, followed by 10–15 fixed/jittered sessions, and then 10–20 short/long sessions. All sessions contained ∼2,000 audiovisual stimuli.

- In random sessions, ISIs were drawn from a uniform distribution (0.5–2.5 s).
- In fixed/jittered sessions, 5 fixed blocks and 5 jittered blocks were presented: blocks of 200 trials alternated between fixed ISIs (1.5 s) and jittered ISIs (uniform distribution: 0.5–2.5 s). Total of 4% of ISIs in fixed/jittered sessions were ‘deviant’, evenly split between short- and long-deviant ISIs.
- In short/long sessions, 19 short blocks and 19 long blocks were presented: blocks of 50 trials alternated between short ISIs (1 s) and long ISIs (2 s).

### Stimulus design

Auditory and visual stimuli were always presented simultaneously and lasted 200 ms on each repetition. The ISI was defined as the absence of the auditory stimulus accompanied by a neutral gray screen.

Auditory stimulus was an 11 kHz pure tone. Visual stimuli were full-field, static sinusoidal gratings of four orientations (0°, 45°, 90°, 135°), at spatial frequency 0.08 cycles/degree. Visual stimuli were displayed on a monitor (Dell, G2524H; 1920x1080 pixels), set to 100% contrast and 75% brightness and covered with a blue screen filter (Amazon) to minimize imaging artifacts during stimulus presentation. The monitor was positioned to the right side of the mouse, with placement parameters following previous literature (*129, 130*).

The same grating orientation was presented for 15-20 consecutive trials before switching to another randomly-selected orientation. Orientation changes did not occur for 3 trials immediately after transitioning to a new block (in both fixed/jittered and short/long sessions) or for 3 trials at the end of each block. Stimulus orientation changes were not analyzed in this study.

### Surgical procedure: window craniotomy

Stereotaxic surgeries (David Kopf, Instruments; Models 942 and 1900) were performed under isoflurane inhalation anesthesia (Somno; Kent Scientific). Anti-inflammatory drug Dexamethasone (1-2mg/kg) was subcutaneously injected 24h and 1h before the surgeries. Post-op care involved subcutaneous injection of the analgesic drug Ketoprofen (2-5 mg/kg). During the surgery, 0.1-0.2mL sterile saline was Intraperitoneally (IP) injected to compensate for liquid loss.

A circular craniotomy in 3mm diameter using a biopsy punch (Fischer Scientific) was made over the left PPC (*89*) (stereotaxic coordinates: 2mm posterior to bregma, 1.7mm lateral) or the left VIS (*131*) (stereotaxic coordinates: 2.92mm posterior to bregma, 2.0mm lateral).

Injection pipettes (15-20 um in diameter; Drummond Micropipettes 3.5”) were back-filled with mineral oil (Sigma Aldrich) and then front-filled with viral vectors: AAV9-Syn-jGCaMP8f (Addgene #162376) for VIP-tdTomato mice, and AAV1-CAG-flex-jGCaMP8f (Addgene #162382) for SST-cre mice. Virus titers was 2.3 x 10^13 gc/mL, and they were both 2-10X diluted in sterile saline. Virus injections were made (Nanoject III; Thomas Scientific) with the speed 10nl/sec in 3-4 different locations (within 1mm around the coordinate center), each with 500nl virus, and ∼150 um below the dura.

A glass plug—made by stacking a 3mm coverslip and a 5 mm coverslip with optical bond (Norland Optical Adhesive NOA 81; Edmund Optics)—was used to cover the craniotomy window. Vetbond (Amazon) and metabond (C&B Metabond® Quick Adhesive Cement System; Parkell Inc.) were used to seal the craniotomy window. Black cement (Ortho-Jet BCA) was applied to reduce possible light leakage during optical imaging.

### Two-photon calcium imaging

Two-photon calcium imaging data were acquired using a Bruker Ultima 2Pplus laser-scanning microscope (Tunable laser set at 920nm: Spectra-Physics Mai Tai High Power DeepSee—Nikon objective: 16X; NA 0.8—Chroma emission filters: 520/40-2p for the green channel and 660/40-2p for the red channel). All imaging experiments were conducted in darkness. The microscope objective was immersed in water-based ultrasonic gel, and the objective was covered with black electrical tape to minimize light leakage.

After surgical recovery, mice were habituated to the setup for 1–2 days before imaging sessions commenced. The same field of view was imaged across days to enable direct comparison of neural responses across the three paradigms. Each imaging session lasted ∼1 hour. During each session, calcium imaging videos (512 × 512 pixels, 30 Hz), stimulus voltage traces (10 kHz), and high-speed videos of the mouse’s left eye were recorded (30 Hz; FLIR camera; Lens: CM3-U3-13Y3M-CS 16mm; each video frame triggered by the imaging scope).

### Preprocessing imaging data

We used a customized pipeline based on suite2p (*132*) to extract neurons and their activity from the imaging videos. Raw videos were imported into suite2p to identify regions of interest (ROIs) and their raw fluorescence traces. All parameters were kept at the software’s default settings.

As an initial quality control step, ROIs with a compactness value greater than 1.06 were excluded to remove poorly segmented ROIs with abnormal geometry. For each trace, a Gaussian filter (*σ* = 600*s*) was applied to estimate the fluorescence baseline and compute 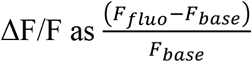, where *F*_*fluo*_ is the raw fluorescence signal and *F*_*base*_ is the extracted baseline from the Gaussian filter. Finally, each neuron’s ΔF/F trace was z-scored individually.

### Identification of VIP interneurons

We used the ‘cellpose’ algorithm (*133*) to identify VIP interneurons labeled with tdTomato in the red channel. For each segmented ROI, we collected the corresponding functional mask from the green channel and quantified red–green co-localization with an overlap score based on intersection-over-union as 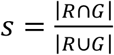, where *R* and *G* are the tdTomato and GCaMP masks, respectively. Neurons were then classified as excitatory (*s* ≤ 0.3), uncertain (0.3 < *s* < 0.7), or VIP (*s* ≥ 0.7). Here, the two thresholds (0.3, 0.7) were chosen a priori based on empirical validation.

### Temporal receptive field (TRF) model

We modeled single-neuron temporal responses around a stimulus using an ex-Gaussian temporal receptive field (TRF) model (**Fig. 1I**; **Fig. 7C,I**; **Fig. S2,3A**). The model has been successfully used to analyze the neural ramping dynamics in the anterior lateral motor cortex (Affan et al., 2024), medial prefrontal cortex (Cao et al., 2024), and entorhinal cortex (Bright et al., 2020). A similar model also has been used to reveal sequential firing in hippocampus and prefrontal cortex (Cruzado et al. 2020).

This model, denoted as *TRF*(*t*) = (*G ∗ E*)(*t*), is defined as

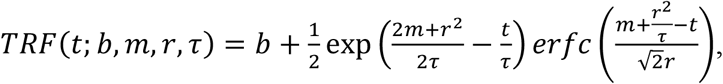

where *erfc* denotes the error function. This form results from convolving a Gaussian kernel *G* :

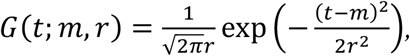

with a causal exponential decay *E* :

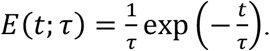

Here, *b* ∈ ℝ is the baseline offset, and *m* ∈ ℝ denotes the response latency (mean of the Gaussian kernel). The parameter *r* ∈ (0,1) controls temporal receptive field width, with smaller values yielding narrower Gaussian kernels. The exponential component is governed by τ > 0, which sets the timescale of ramp-up or ramp-down. Responses with larger τ decay more slowly back to baseline. For interpretability, we use 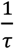 to represent ramping speed.

To fit the TRF model to a neuron (**Fig. 1I**), we aligned the neural activity during the interval before and the one after stimulus onset at *t* = 0 . Time was normalized for fitting stability, denoted as 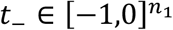 and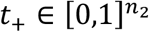, respectively, where *n*_1_ and *n*_2_ are the number of time points.

Let 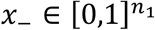 and 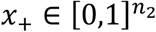be the two segmentations of the normalized activity, referring to the pre-stimulus and post-stimulus activity, respectively. We then fit *TRF*(−*t*; *b, m, r*, τ) to *x*_ and fit *TRF*(*t*; *b, m, r*, τ) to *x*_+_, which we refer to as ramp-up and ramp-down models, respectively (**Fig. 1I**).

Note that both models have the same parameters, but they are fit to inputs of opposite direction (flipped in time). The ramp-up and ramp-down models differ only by the direction of the time input (forward vs reversed). This gives two sets of parameters *θ*___ = {*b*___, *n*___, *r*___, τ___} and *θ*_+_ = {*b*_+_, *n*_+_, *r*_+_, τ_+_}, referring to the *ramp-up* and the *ramp-down* model components.

Note that this model always assumes that each neuron has both the ramp-up component, before stimulus, and ramp-down component, after stimulus. The goodness of fit is evaluated by the coefficient of determination between the prediction and the ground-truth normalized activity for both components.

### Clustering neural responses

We used a combination of generalized linear modeling (GLM) (*42, 134*) and the TRF model prior to clustering (**Fig. 1I**). For each paradigm, neural responses from all sessions were pooled together, and the following steps were performed on the combined dataset.

1. **GLM modeling**. High-sampling-rate voltage recordings of the stimulus trace (10 kHz) were first aligned with the fluorescence traces, for the entire 1hr session, using nearest-neighbor interpolation. For each neuron, we fit a GLM to the z-scored ΔF/F trace 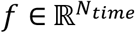 using the binary stimulus trace 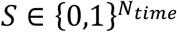 as input, over the window [−1500,1500] ms relative to stimulus onset (*N*_*time*_ denotes the number of imaging frames in a session, with 1 indicating stimulus presentation and 0 indicating no stimulus). This window was determined such that we can fit the TRF model easily for the incoming steps. To prevent overfitting and obtain smooth kernel estimates, we applied L2 regularization *α* = 5 × 10^4^ multiplier). The GLM kernel for the i-th neuron 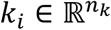 was then estimated via the least square principle *k*_*i*_ = (*S*^*T*^ *S*)^-1^*S*^*T*^*f*. Next, since the fit convolution kernel is flipped from the actual GLM kernel, we flipped the resulting GLM kernel to correct this. We then split the fit GLM kernel *k*_*i*_ into two parts, *k*_*i*_ (*t*) with *t* < 0 and *k*_*i*+_ (*t*) with *t* > 0, which are therefore the summaries of how a neuron generally responds before and after a stimulus, respectively.
2. **TRF modeling**. See the Methods section ‘Temporal receptive field (TRF) model’ for details. In brief, for each neuron, we fit 2 TRF models to the 2 components of the GLM kernel, *k*_*i*__ (*t*) *and k*_*i* +_(*t*), corresponding to the ramp-up and ramp-down models, respectively:

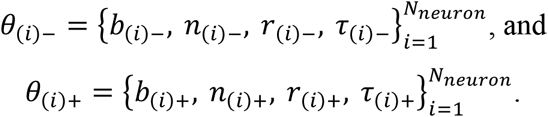

Let 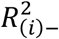 and 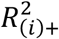 denote the goodness of fit for the two models. We defined ramp-up neurons as the neurons that can fit (1) well with the ramp-up model, and (2) better than the ramp-down model, and vice versa. Let *θ* be the threshold (*θ* = 0.4 in this paper). Thus, those neurons with both 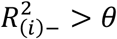 and 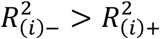 are determined as the ramp-up neuron, while those neurons with both 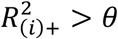 and 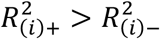 are the ramp-down neuron. All remaining neurons with goodness-of-fit values below the threshold for both categories were classified as the “mixed” cluster (**Fig. 2E,K; Fig. S3; Table 2,3**), and excluded from the main figures. This results in *N*_*up*_ ramp-up neurons and *N*_*down*_ ramp-down neurons.
3. **Hierarchical clustering**. After getting the two board categories of neurons, within each ramp-up and ramp-down group, we use hierarchical clustering on the fit the temporal receptive field width 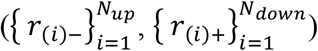, and rising time constant 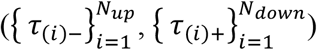, to find subgroup of neurons with distinct response dynamics. The number of clusters is determined such that the Silhouette Coefficient is maximized.

### Temporal scaling analysis

To test for potential temporal scaling effects (**Fig. 3C,F**), we performed the following procedures. Let 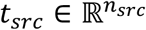 denote the original time stamps for an aligned neural trace and 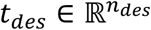 denote the target time range to which the trace was scaled, where *n*_*src*_ and *n*_*des*_ are the respective length of the time stamps. Scaled time stamps were computed as:

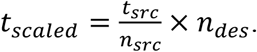

A nearest neighbor interpolation was then used to map *t*_*scaled*_ to *t*_*des*_. This procedure left the neural activity values unchanged but adjusted the time axis to match the target interval. Given neural activity aligned to stimulus onset, we applied this scaling separately to the pre-stimulus and post-stimulus intervals, while leaving the activity during the stimulus itself unchanged.

### Modulation index for deviant ISIs in fixed/jittered sessions

We quantified event-evoked response changes (**Fig. S4A**) with a modulation index (MI) defined as

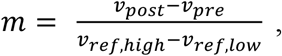

where 𝒱_*pre*_ and 𝒱_*post*_ are the average signal within the pre- and post-event windows (see below), and 𝒱_*ref,high*_ and 𝒱_*ref,low*_ are the highest and lowest 10% activity values in a reference window ([-2.5, 4.0] s from the deviant in all cases).

The difference between post- and pre-event responses was normalized by the neuron’s reference activity range, and treats increases and decreases symmetrically (*m* > 0: relative increase after the event; *m* < 0: decrease; and *m* = 0 no change). Compared with the conventional definition of modulation index (difference divided by sum), it normalizes the post–pre difference by an independent reference activity range, avoiding unstable values given small denominators. It also reduces saturation near the bounds while keeping most values within [-1,1], allowing strong modulations to remain more distinguishable.

For the unexpected omission during long deviant ISIs (negative prediction errors), modulation index reflects omission response modulation relative to baseline immediately before the omission (**Fig. S4A**, left). 𝒱_*pre*_ and 𝒱_*post*_ was computed from the average activity in a window of size 0.5 s immediately *before* (*after*) the expected stimulus time.

For the stimulus immediately after deviant ISIs (positive prediction errors), 𝒱_*pre*_ and 𝒱_*post*_ were computed by averaging activity values above the 30th percentile within the [-0.1, 0.5] s window relative to stimulus onset. 𝒱_*pre*_, was aligned to the pre-deviant stimulus, 𝒱_*post*_ to the post-deviant stimulus (**Fig. S4A**, middle and right); hence, the modulation index reflects response modulation relative to the ‘standard’ stimulus.

Two versions of the MI were computed: single-trial, neuron-averaged in **Fig. S4**, and single-neuron, trial-averaged in **Fig. S6**. For both versions, the same quantification windows, described above, were used. In **Fig. S6**, only neurons with modulation indices above 0.1 or below -0.1 were selected.

### Population trajectory analysis

Neural population activity was analyzed in a low-dimensional latent space using principal component analysis (PCA). For each condition c_*i*_ (e.g., trials within the fixed block or jittered block in **Fig. 5D,H,M,Q**, or trials within each ISI bin in **Fig. 6H,P**), we first averaged neural activity across trials to obtain a condition-specific response matrix: 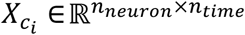.

These matrices were then concatenated across conditions to form a combined data matrix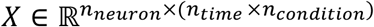. PCA was fitted to this combined dataset to extract the principal axes capturing the greatest shared variance across neurons and conditions.

For visualization, population trajectories for each condition were obtained by projecting their time-varying population activity onto the top three principal components (PC1–PC3), which together defined a shared low-dimensional space for comparing neural dynamics across conditions. Each point along a trajectory thus represents the instantaneous population state at a given time, and its color encodes time progression. To obtain smoother neural trajectories, each PC was interpolated to increase the number of time points fourfold, followed by Savitzky–Golay filtering (polynomial order = 3, window size = 8).

Population dimensionality was also estimated using PCA (**Fig. 2F,L**). Data from the long block of short/long sessions were used for this analysis. Neural activity was first averaged across trials within the long-ISI block to obtain a population activity matrix 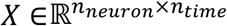 . PCA was then applied separately to the population activity matrix for each cluster. We quantified the dimensionality of each cluster by extracting the cumulative explained variance across principal components, which reflects how many components were required to capture the structure of population activity.

### Temporal decoding analysis

We used the long block of the short/long sessions for this analysis. To evaluate how neural populations encode elapsed time (**Fig. 7**), we trained classifiers to discriminate neural activity between pairs of time bins within an interval. Neural data were aligned to stimulus onset and truncated at the onset of the next stimulus. Each time bin was defined as two non-overlapping imaging frames, with neural activity averaged within each bin for every neuron and trial.

To reduce single-trial noise in calcium imaging, we used a bootstrapping approach: on each iteration (n=250), pseudo-trials were generated by averaging random subsets of three trials. The resulting data tensor was denoted as: 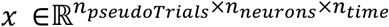, where *n*_*pseudoTrials*_, *n*_*neurons*_, *n*_*times*_ are the number of pseudo trials, neurons, and time bins, respectively.

PCA was then used to reduce *n*_*neurons*_ to 50 dimensions (d), preserving >0.99 cumulative explained variance (**Fig. 2F,L**). Using the same number of components across clusters enabled comparison of decoder performance between clusters containing different numbers of neurons.

We used stratified randomized cross-validation. For each pair of time bins, pseudo-trials were randomly divided into training and test sets using an 80/20 split. A Support Vector Machine (SVM) classifier with a linear kernel was trained on data shaped as *d* × *n*_*pseudoTrials*_ × *n*_*time*_ and evaluated on the held-out test set (d: top 50 PCA components). This train/test procedure was repeated 25 times using stratified randomized folds across all time-bin pairs, and the results were assembled into a time-by-time accuracy matrix: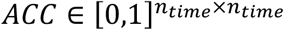.

### Cell-type interaction analysis

We used Spearman correlation coefficients as a proxy for interaction strength within and across simultaneously imaged VIP and excitatory populations. Spearman correlations were chosen to minimize the influence of activity magnitude on correlation estimates.

During fixed and jittered sessions, single-trial neural activity was aligned to the stimulus before the deviant ISI. For each neuron pair, the correlation was computed along the trial dimension at each time point, using trial-by-trial neural responses. This procedure generated a time-resolved correlation trace for each neuron pair. Correlation values were then averaged across pooled neuron pairs from all recording sessions to obtain the mean population-level correlation over time.

## Acknowledgments

We are grateful to the Haider Lab for providing the SST-Cre mice, to Dieter Jaeger, Simone Russo, and Corbett Bennett for thoughtful discussions, to Laurence Copeland and Vanshika Mehta for assistance with early experiments, to Joey Broussard, Robert Howard, Leslie Claar, and Ben Ouellette for valuable technical expertise.

## Funding

This work was supported by:

Whitehall Foundation

Research Corporation for Science Advancement

Chan Zuckerberg Initiative

Georgia Institute of Technology startup funds

## Author Contributions

Conceptualization, Writing: F.N., Y.H.

Data Analysis: Y.H.

Engineering Support: T.S.

Experiments: Y.H., M.C., S.W., S.M.

Supervision, Funding Acquisition, Resources: F.N.

Visualization: Y.H., A.S., S.M.

## Data, Codes, and Materials Availability

All data used in this paper will be made public upon the paper publication. Code for data processing and analysis is publicly available on Najafi Lab GitHub page: https://github.com/najafi-laboratory/2p_imaging/tree/main/passive_interval_oddball_202412

## Supplementary Materials

**Fig. S1.**
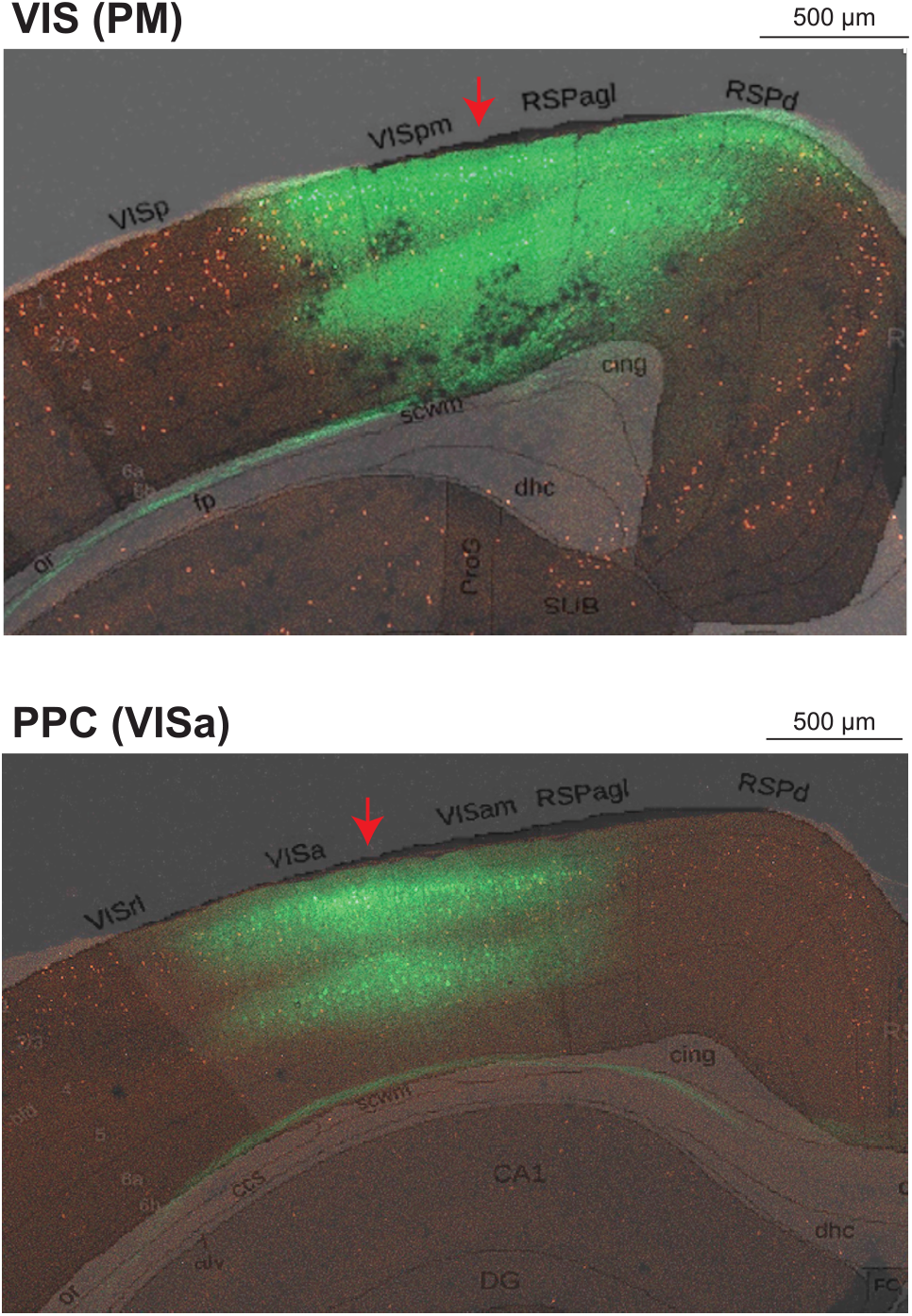
Histological verification of imaging sites in the medial visual and posterior parietal cortex. Histology images from two example mice (VIP-tdTomato) imaged in VIS (**top**, area PM) or PPC (**bottom**), with Allen coronal slices superimposed. Red arrow: center of craniotomy window used for imaging, marking VIS (PM), and PPC (VISa). Green: cells labeled by GCaMP8f injection. Red: VIP neurons. Multiple injections were performed within the 3-mm cranial window. Two-photon imaging was primarily conducted from the center of the imaging window.

**Fig. S2.**
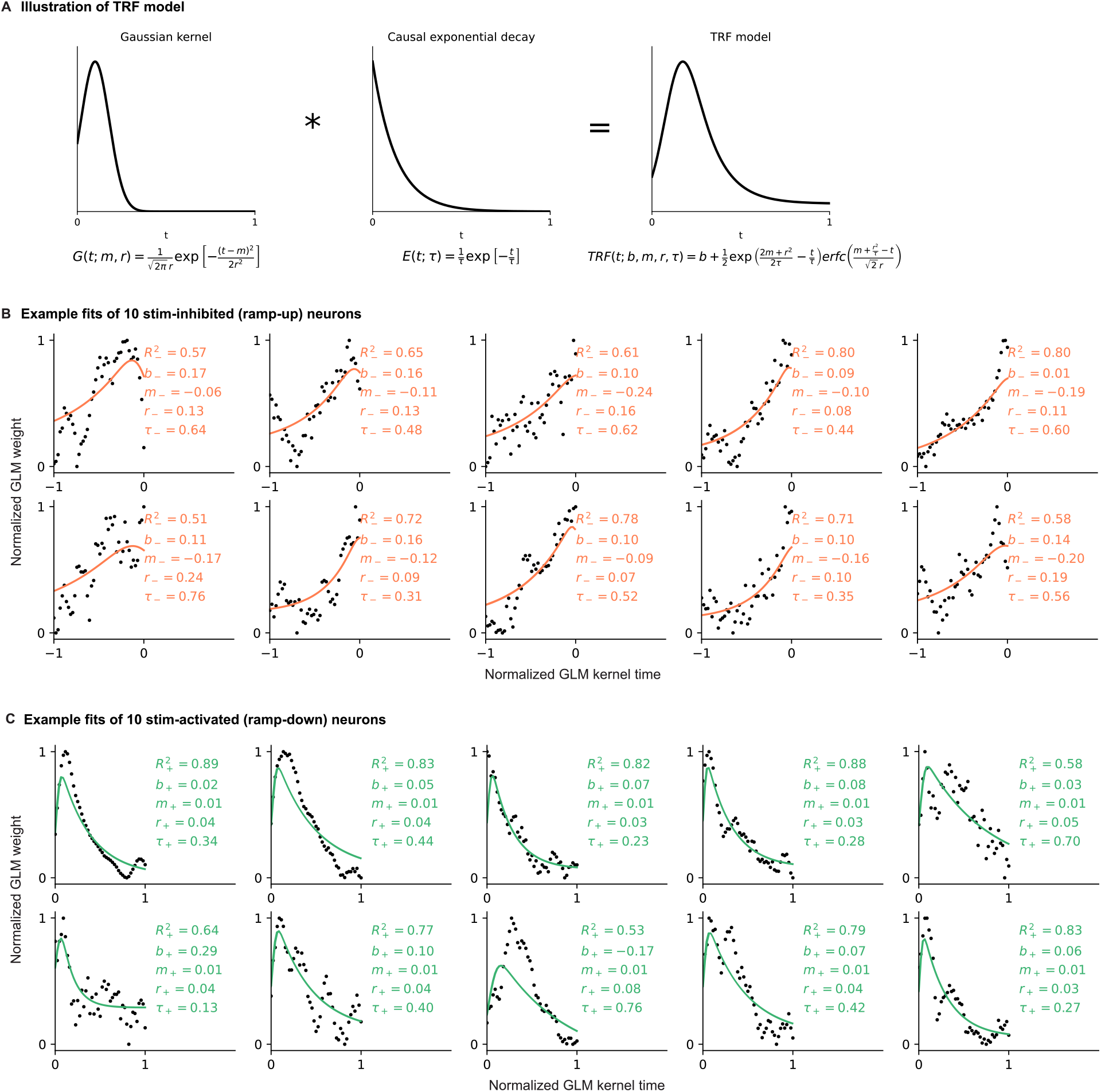
Example fits of GLM kernels using the TRF model. In all panels, legends report goodness-of-fit (R^2^), and model parameters: baseline (b), latency (m), temporal width (r), and time constant (τ), for each ramp-up (-) and ramp-down (+) component. **(A)** TRF model function is the convolution results between a Gaussian kernel and a causal exponential function. **(B)** Ten examples of GLM kernel components fit by the ramp-up model. Goodness-of-fit values and model parameters are indicated. Dots: normalized GLM kernel weights before time 0. Green: ramp-up TRF model fit. **(C)** Same as **(A**), but for components fit by the ramp-down model. Orange: ramp-down TRF model fit.

**Fig. S3.**
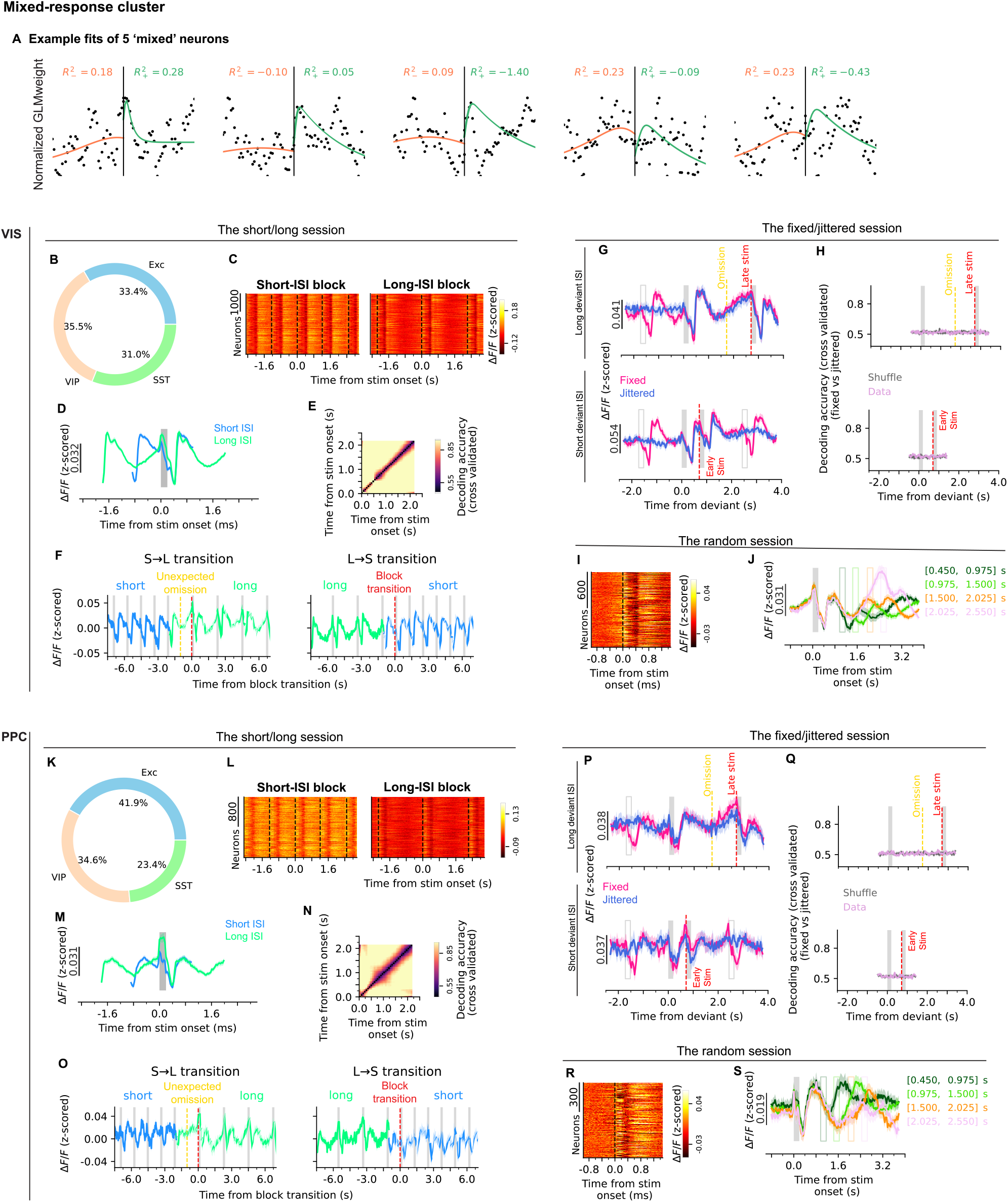
Neurons in the “mixed” cluster reproduce the main findings across all three paradigms. Activity of neurons lacking exclusively ramp-up or ramp-down dynamics, and therefore excluded from the main figures, is shown across all three paradigms. **VIS:** top panels; **PPC:** bottom panels. **(A)** Five examples of GLM kernel components fit by the ramp-up (green) and ramp-down (orange) models. Goodness-of-fit values for both models is indicated. These neurons are grouped in the ‘mixed’ cluster because their goodness-of-fit for neither the ramp-up or down models exceeded our threshold (0.4) determined empirically. Dots: normalized GLM kernel weights centered at time 0 (black vertical line). **(B-F)** Data from short/long-interval sessions, demonstrating rapid response changes immediately following block switches. **(B)** Fraction of cell types within the “mixed” cluster. **(C)** Heatmaps of trial-averaged, single-neuron activity aligned on stimuli (vertical dashed lines) in the short (Left) and long (Right) ISI blocks. Similar to **Fig. 3A**. **(D)** Neuron- and trial-averaged activity for short (blue) and long (green) blocks, mean ± 95% confidence interval, aligned on the stimulus (gray box). Similar to **Fig. 3B**. **(E)** Pairwise classification accuracy matrices (cross-validated; linear SVM). Each cell shows the accuracy for distinguishing a given pair of time bins. Similar to **Fig. 7B**. **(F)** Trial-by-trial neural activity around block transitions, mean ± 95% confidence interval, aligned on the block transition. Similar to **Fig. 4B**. **(G-H)** Data from fixed/jittered-interval sessions, demonstrating similar deviant responses—to omissions and early/late stimuli—across fixed and jittered blocks. **(G)** Neural responses (mean ± 95% confidence interval) for each cluster, for fixed (red) and jittered (blue) blocks, aligned on the stimulus before the deviant ISI (Top: long deviant ISI; Bottom: short deviant ISI). Similar to **Fig. 5A,E**. **(H)** Decoding accuracy for classifying fixed vs. jittered trials (cross-validated; linear SVM; Purple: real data; Gray: shuffled data; Yellow line: omission; Top: long deviant ISI; Bottom: short deviant ISI). Similar to **Fig. 5B,F**. **(I-J)** Data from the random-interval sessions, demonstrating the evolution of responses during the ISI. **(I)** Heatmap of trial-averaged, single-neuron activity aligned on stimulus onset (vertical dashed line). Similar to **Fig. 6A**. **(J)** Random ISIs divided into four bins (different colors). Neural responses averaged across trials within each ISI bin. Similar to **Fig. 6F**. **(K-S)** Same as **(B-J)**, but for PPC.

**Fig. S4.**
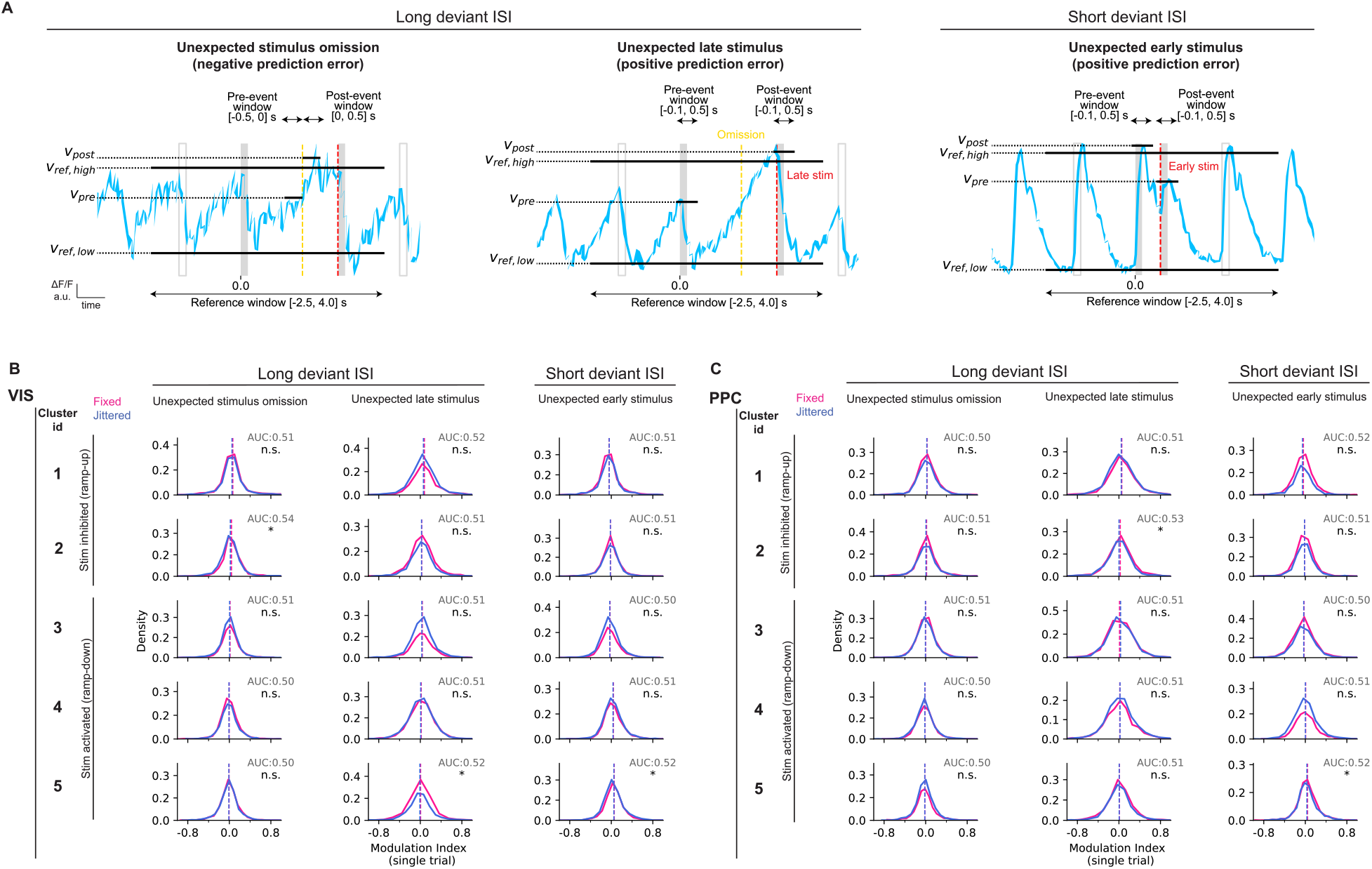
Responses to temporal deviations are similar in fixed and jittered contexts, arguing against temporal predictive processing as the underlying mechanism of these neuronal dynamics. **(A)** Illustration of the modified modulation index with an example. It is computed as (𝒱_*post*_ – 𝒱_*pre*_)/(𝒱_*ref,high*_ – 𝒱_*ref,low*_). Solid horizontal lines specify quantification windows. 𝒱_*ref,high*_ and 𝒱_*ref,low*_ are computed from the highest and lowest 10% activity values within the same reference window, respectively. The pre- and post-event windows are distinctly defined for each case (omissions, late stimuli, early stimuli), but the reference windows are the same for all cases. **(B)** Modulation indices were computed on single-trial, neuron-averaged responses for each cluster (top to bottom), separately for fixed and jittered trials. Distribution of modulation indices across fixed (pink) and jittered (blue) trials (dashed line: distribution mean). ROC analysis was performed to compare the two distributions. Area under the curve (AUC) values are reported, along with statistical significance (n.s.: non-significant; asterisk: p < 0.05). Statistical significance was assessed using a two-sided permutation test (1000 permutations), in which trial labels were randomly shuffled to generate a null distribution of AUC values centered at chance (0.5). **(C)** Same as **(B)**, but for PPC.

**Fig. S5.**
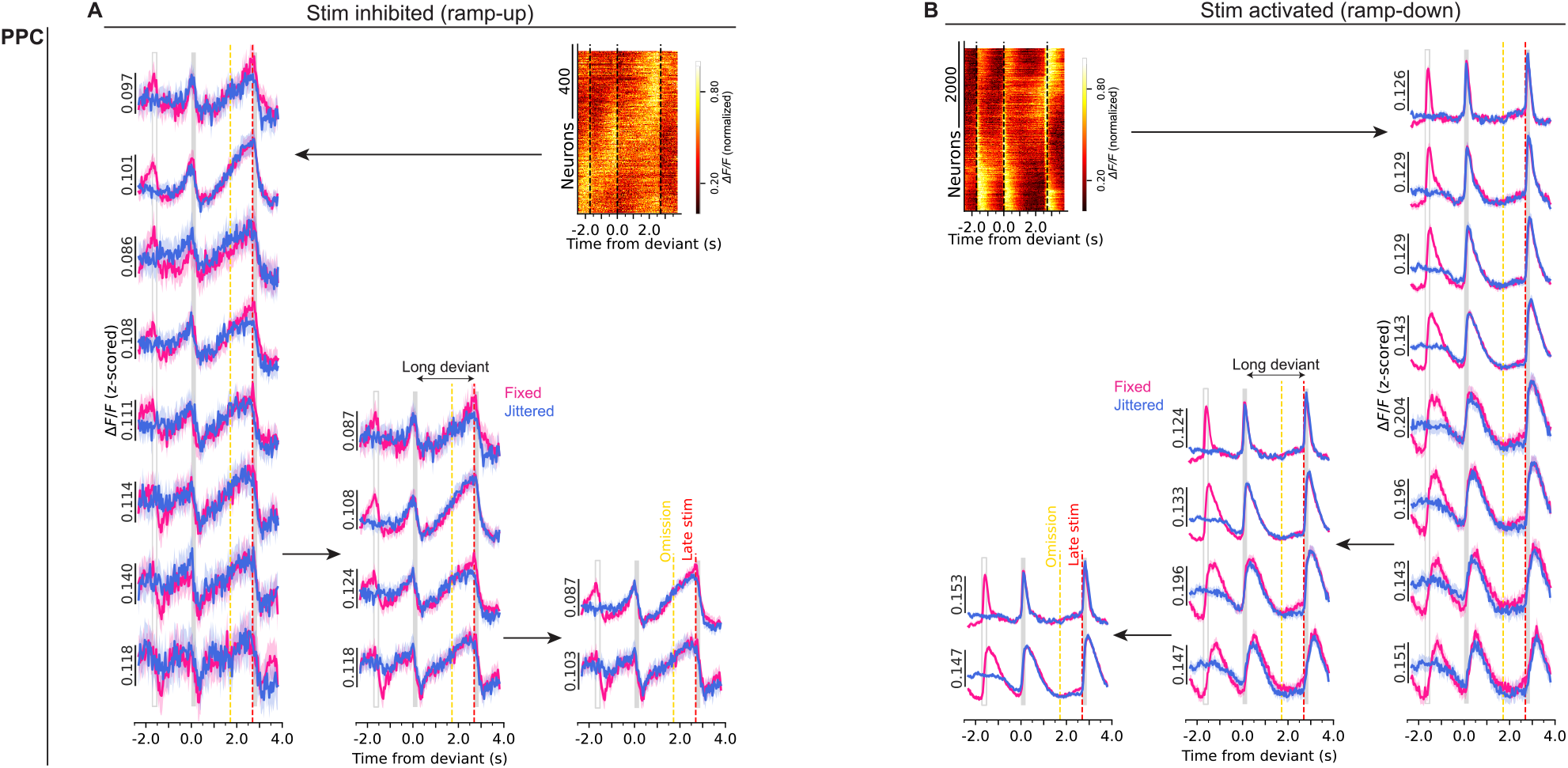
Intermediate hierarchical subclusters also do not exhibit differential responses between fixed and jittered ISI blocks during long-deviant trials. Data from PPC. Fixed/jittered-session data during long-deviant ISI trials. Intermediate subclusters are shown throughout the hierarchical merging process for stimulus-inhibited (**A**) and stimulus-activated (**B**) neurons. Heatmaps show single-neuron trial-averaged activity during fixed long-deviant trials (dashed lines: stimulus onset), sorted using Rastermap (*135*). Arrows indicate the hierarchical merging process from individual neurons to 8, 4, and 2 clusters. Neural responses (mean ± 95% confidence interval) for each cluster are shown for fixed (red) and jittered (blue) blocks, similar to **Fig. 5A,J**.

**Fig. S6.**
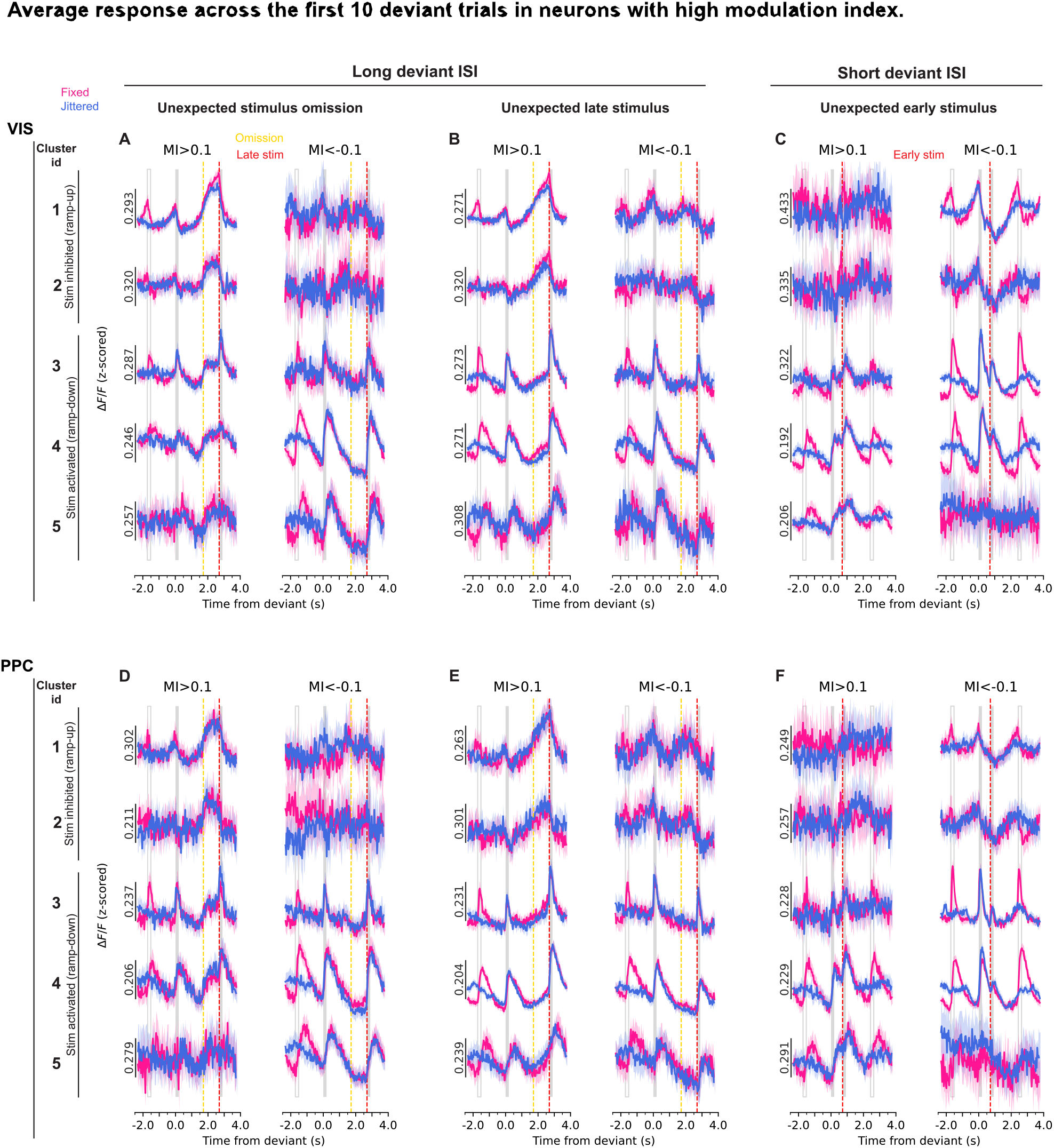
Selected neurons with high modulation indices still do not exhibit differential responses between fixed and jittered ISI blocks during early deviant trials. **VIS:** top panels; **PPC:** bottom panels. Modulation index was computed on trial-averaged activity for individual neurons using quantification windows shown in **Fig. S4A**. Neurons with large modulation indices (above or below 0.1) were selected, and their average activity across the first 10 deviant trials is shown, aligned on omissions during long deviant ISIs **(A)**; late stimuli after long-deviant ISIs **(B)**, and early stimuli after short-deviant ISIs **(C). Left**: neurons with modulation index < −0.1. **Right**: neurons with modulation index > 0.1. **(D-F)** Same as **(A-C)**, but for PPC.

**Fig. S7.**
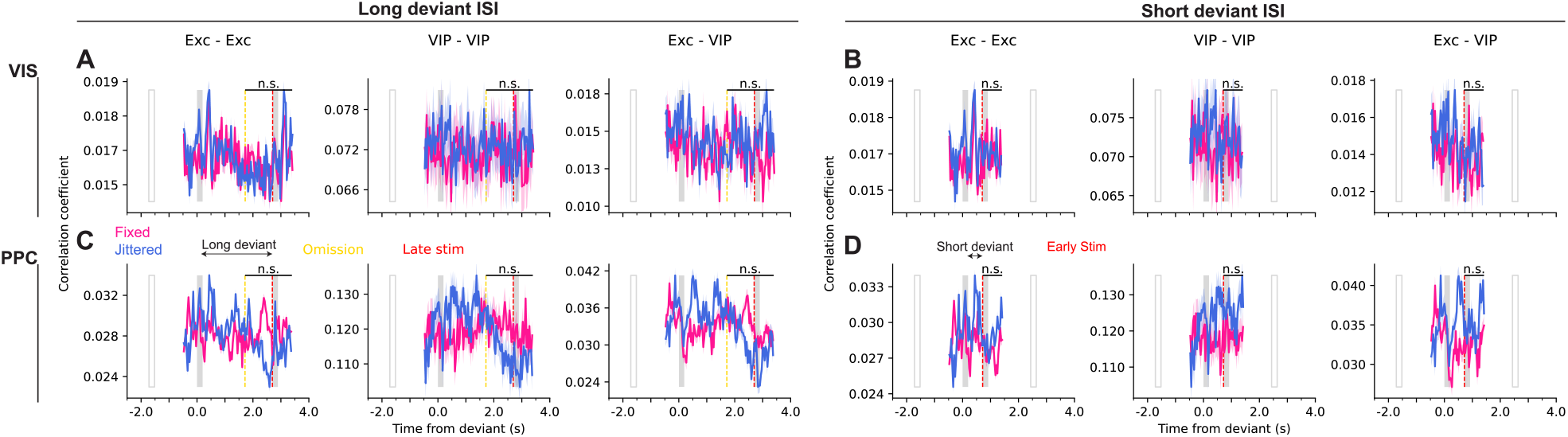
VIP and excitatory interactions during deviant intervals. **VIS:** top panels; **PPC:** bottom panels. Spearman correlation coefficients between simultaneously-imaged excitatory and VIP neurons during long **(A)** and short **(B)** deviant intervals, in fixed/jittered sessions. Traces are aligned on the stimulus before the deviant ISI. For each session, correlation was computed between pairs of neurons across trials, separately for each timepoint within the trial. **Exc-Exc:** correlation within excitatory population. **VIP-VIP:** correlation within VIP population. **Exc-VIP:** correlation between excitatory and VIP populations. Mean ± 95% confidence interval, pooled across all neuron pairs, from all sessions and all mice. Yellow line: expected but omitted stimulus; Red: unexpected late or early stimulus; Gray boxes: stimuli; Empty boxes: stimuli in fixed and mean stimuli time in jittered). Black lines: statistical test window (n.s.: non-significant; Wilcoxon signed-rank test). **(C-D)** Same as **(A-B)**, but for PPC.

**Fig. S8.**
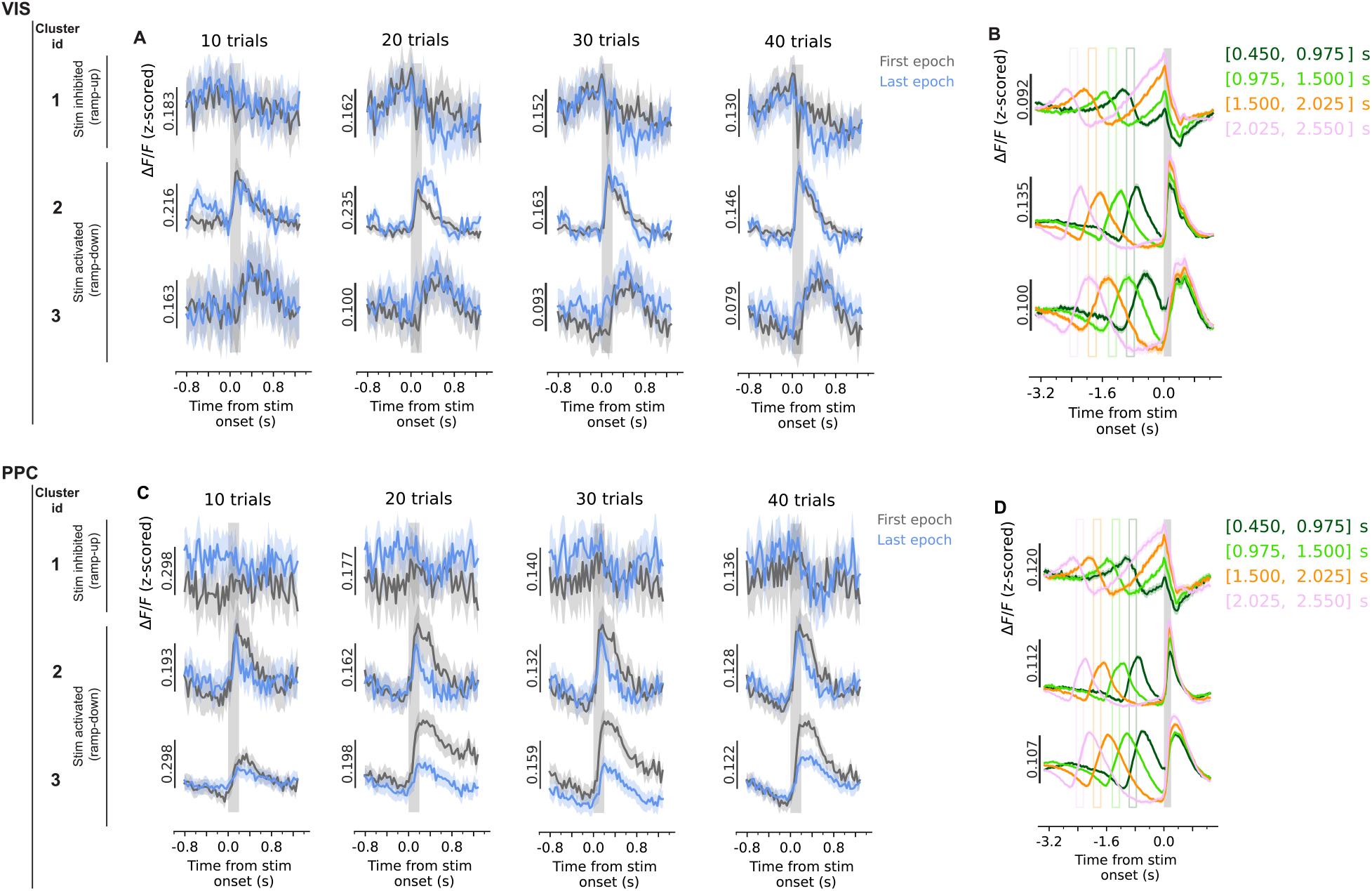
Responses across the initial trials of the random session confirm that extensive exposure is not required for pre-stimulus ramping activity to emerge. Averaged responses are shown for the first epoch (gray) and last epoch (blue) of the first session. Epochs were defined using different trial numbers to compare response profiles across the beginning and end of the session. Pre-stimulus ramping is evident in VIS in the first 20 trials. Data are noisier in PPC, due to its fewer number of ramp-up neurons (**Fig. 6O**). **VIS:** top panels; **PPC:** bottom panels. **(A)** Left to right: averaged responses for the first (gray) and last (blue) 10, 20, 30, 40 trials of the first random session, respectively, similar to **Fig. 6D**. **(B)** Random ISIs divided into four bins (different colors). Neural responses averaged across trials within each ISI bin, and aligned to the subsequent stimulus, illustrating larger stimulus-evoked responses following longer preceding ISIs. Gray box: stimulus. Empty boxes: mean ISI of each bin. **(C-D)** Same as **(A-B)** but for PPC.

**Fig. S9.**
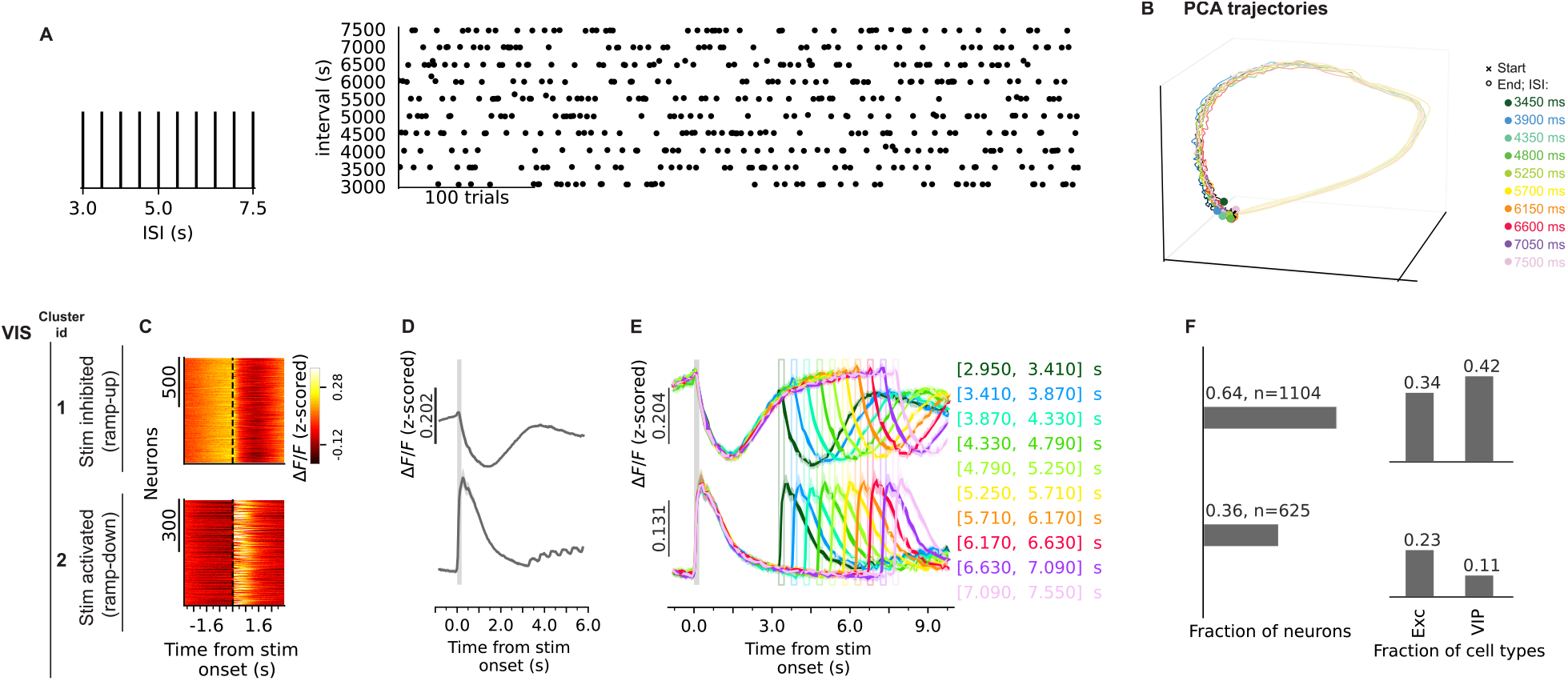
Ramp-up neural responses plateau about 4s after stimulus onset. Data from extended-ISI random sessions (area VIS; 3 VIP-tdTomato mice; 5 sessions per mouse). **(A)** The extended random session is similar to the random session in **Fig. 1H** and **Fig. 6**, but with discrete, uniformly distributed ISIs in the range 3.0–7.5 s. **Left:** distribution of ISIs. **Right:** example sequence of consecutive trial ISIs. **(B)** Principal component analysis (top three components) of population activity from stimulus onset through the ISI. Trajectories follow a common manifold across intervals, extending further with longer ISIs. **(C)** Heatmaps of trial-averaged, single-neuron activity aligned on stimulus onset (black line), for each cluster (top to bottom). **(D)** Neuron- and trial-averaged responses aligned on the stimulus (gray box) for each cluster. **(E)** Neural responses averaged across trials within each ISI (different colors). Gray box: stimulus; vertical lines: ISIs. **(F) Left:** fraction of all neurons assigned to each cluster. **Right:** fraction of neurons of each cell type assigned to each cluster.

**Fig. S10.**
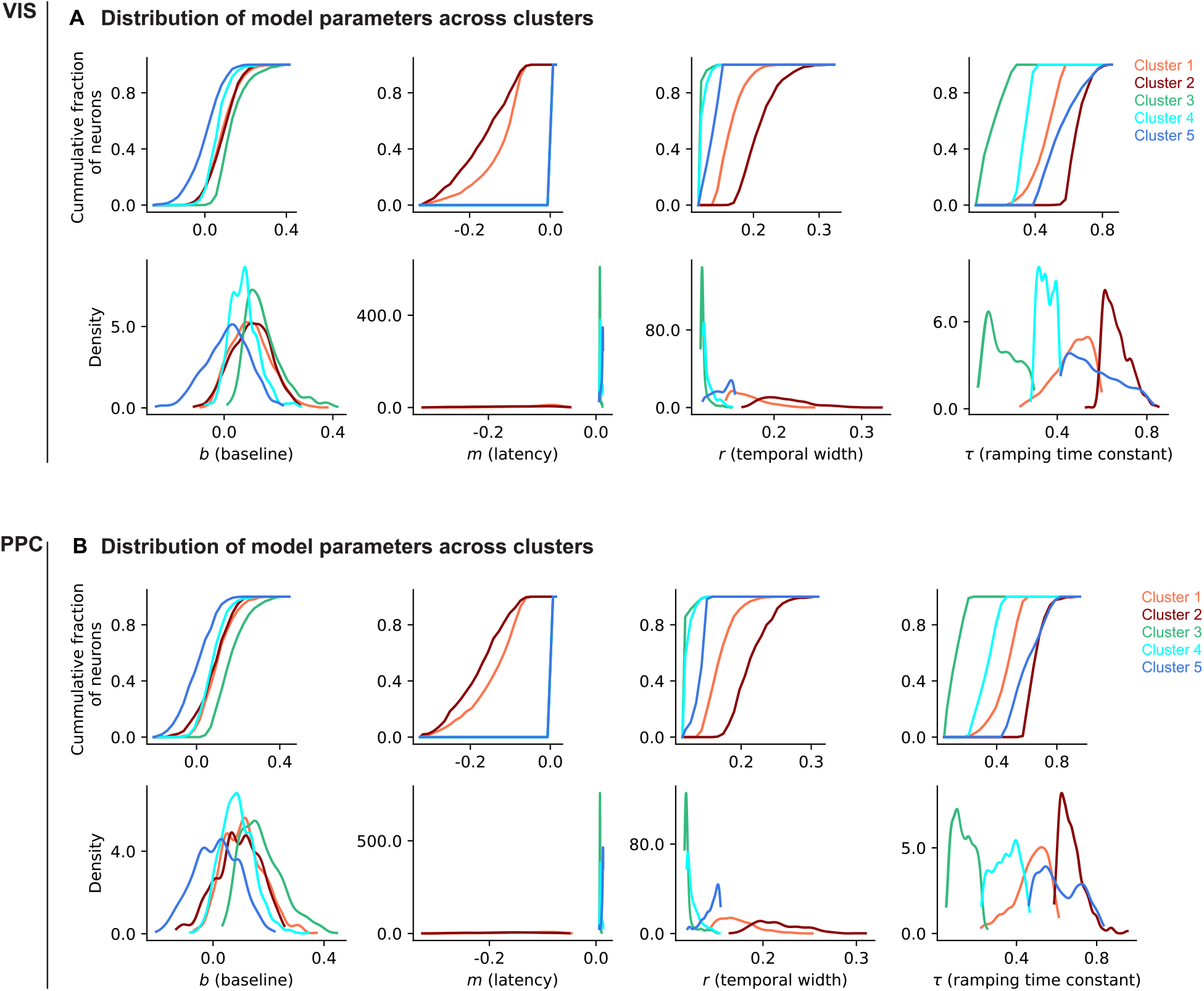
Distribution of TRF model parameters. Data from the long block of the short/long sessions. Comparison of the 4 parameters of the temporal-receptive-field (TRF) model across all clusters (Orange, red: ramp-up clusters; Green, cyan, blue: ramp-down clusters). **VIS:** top panels; **PPC:** bottom panels. **Top:** cumulative histograms. **Bottom:** distribution densities. ‘b’ reflects the baseline shift. ‘m’ reflects the time latency of activity peak, relative to stimulus onset. ‘τ’ is the time constant, reflecting the inverse of ramping speed. ‘r’ reflects the temporal width of the response.

